# Implication of the amyloid precursor protein expressed in GABAergic interneurons for CA3 neuronal dynamics

**DOI:** 10.64898/2026.11.17.767235

**Authors:** Anaël Erhardt, Ana Moreira-de-Sá, Séverine Deforges, Eva Ferreira Rodrigues, Ashley Kees, Christophe Mulle

## Abstract

The amyloid precursor protein (APP), a key player in Alzheimer’s disease (AD), is ubiquitously expressed in a large majority of neurons, including GABAergic interneurons (INTs) in the hippocampus. INT dysfunction and subsequent abnormalities in neural circuits have emerged as potential mechanisms of cognitive decline in AD. How APP expressed in selective classes of hippocampal INTs controls local neuronal activity and thereby influences brain rhythms remains unclear. We investigated the physiological function of APP in GABAergic INTs with regard to intrinsic excitability, synaptic properties and circuit activity in the hippocampal CA3 region. We selectively deleted APP, and its homolog APLP2 to prevent compensatory mechanisms, in CA3 INTs. We categorized INTs into different clusters based on morphological and electrophysiological properties, and detected only subtle differences in intrinsic excitability and dendritic complexity between control and conditional KO INTs. In contrast, the selective deletion of APP/APLP2 in CA3 INTs impaired the amplitude and kinetics of spontaneous inhibitory currents in CA3 pyramidal cells (PCs). We further performed *in vivo* silicon probe recordings within CA3 in head-fixed awake mice to investigate the impact of APP/APLP2 deletion on CA3 circuit activity. By analysing oscillations and the spiking activity of putative CA3 PCs and INTs we revealed alterations in the cross-frequency coupling at oscillatory frequencies in the low gamma range, as well as in the strength of connections specifically between INTs. Our work helps better understand the physiological contribution of APP to network activity and to excitatory/inhibitory balance within hippocampal circuits, which are impaired in AD.

## Introduction

The amyloid precursor protein (APP), alongside the amyloid precursor-like proteins 1 and 2 (APLP1/APLP2), belongs to a well-conserved gene family in the mammalian brain. APP and APLPs are single type I transmembrane proteins with a large N-terminal extracellular domain and a short C-terminal cytosolic tail, being expressed in both excitatory and inhibitory neurons. APP, which undergoes complex proteolytic processing that gives rise to various derivative fragments, is intrinsically linked to the pathophysiology of Alzheimer’s disease (AD). Indeed, a causal relationship exists between mutations of APP (and/or presenilin 1 and 2, which cleaves APP) and autosomal dominant familial forms of AD (Bateman et al., 2011). Abnormal amyloidogenic APP cleavage is associated with cellular and synaptic dysfunction (Palop & Mucke, 2010; Selkoe, 2002). At a circuit level, AD pathophysiology is characterized by alterations of brain oscillations and abnormal network synchrony affecting the communication between brain regions (Busche et al., 2015; Busche & Konnerth, 2016; Palop & Mucke, 2016; Viana da Silva et al., 2023). Furthermore, many observations from animal models of AD and from clinical studies have pointed to deficits of GABAergic interneurons (INTs) with reduced inhibition in brain circuits (Hijazi et al., 2023; Palop et al., 2007; Palop & Mucke, 2016; Xu et al., 2020), associated with network hyperactivity and cognitive dysfunction (Jiménez-Balado & Eich, 2021; Schmid et al., 2016; Verret et al., 2012; Zott et al., 2019). The abnormal hyperactivation of hippocampal neurons is most likely caused by a shift in excitation/inhibition balance (Burns et al., 2026; Palop & Mucke, 2016). Interestingly, there is a reduction of inhibitory densities in brain regions where the pathophysiology has settled in AD patients (Kurucu et al., 2022). The mechanisms by which deficits in the physiology of GABAergic INTs contribute to network and cognitive dysfunctions in AD and related conditions remain to be fully elucidated. Understanding the physiological function played by APP in GABAergic INTs may provide insights into these mechanisms.

GABAergic INTs which express high levels of APP (B. Wang et al., 2014), play a major role in maintaining a proper level of neuronal activity in hippocampal circuits through adaptive mechanisms (Atallah & Scanziani, 2009; Sukenik et al., 2021). GABAergic INTs are heterogeneously distributed within hippocampal circuits and display vast morpho-functional diversity (Pelkey et al., 2017). Interestingly, hippocampal oscillations, local disinhibition and precise spatiotemporal control of principal cells through inhibitory synaptic inputs rely on morphologically and functionally specialized populations of GABAergic INTs (Booker & Vida, 2018; Buhl et al., 1994; Chamberland et al., 2024; Gloveli et al., 2005; Klausberger & Somogyi, 2008).

In pathological conditions, network alterations are potentially caused by the impaired activity of specific GABAergic subpopulations such as parvalbumin-containing (PV) INTs, a subset of fast-spiking inhibitory INTs, predominantly contributing to the generation of gamma oscillations (Hijazi et al., 2023). In knock-in mice overexpressing human mutant APP, PV INTs have reduced levels of voltage-gated sodium channels which lead to aberrant gamma oscillations and network hypersynchrony (Verret et al., 2012). Strikingly, impaired slow-gamma oscillations in AD transgenic mice can be restored *in vivo* by pharmacologically increasing GABAergic synaptic inhibition (Busche et al., 2015). In contrast, in the APP/PS1 mouse model of AD, in which mutant APP is overexpressed, PV INTs were shown to be hyperexcitable at 16 weeks of age, and chemogenetic inhibition of PV INTs to control levels rescues memory deficits (Hijazi et al., 2020). Altered theta-gamma coupling and deficits in PV INT-to-PC synapses in CA1 caused by the presence of pathogenic Aβ oligomers are recovered by light-induced activation of PV and somatostatin-containing (SST) INTs (Park et al., 2020). The tight relationship between AD pathophysiology and neuronal inhibition raises a special interest in studying the physiological role of APP in GABAergic INTs. Direct investigation of the physiological role of APP family members in the hippocampus (HPC) has relied on knock-out strategies in mice, with either ubiquitous deletion or selective suppression of APP in targeted neuronal populations (Gabriele et al., 2022; Müller et al., 2017). Full constitutive APP knock-out (KO) and conditional KO mouse models have displayed alterations in both excitatory and inhibitory synaptic transmission as well as in hippocampal function (Hick et al., 2015; Seabrook et al., 1999). The conditional removal of APP and APLP2 from all GABAergic forebrain INTs disrupts hippocampal plasticity, dendritic branching of CA1 PCs and INTs, and learning and memory capabilities (Mehr et al., 2020). However, the selective role of APP expressed in GABAergic INTs in governing neuronal activity and oscillations within local hippocampal circuits is currently poorly understood.

Here, we aimed at exploring the precise physiological role of APP expressed in GABAergic INTs on local circuit activity in CA3, a brain region particularly vulnerable in AD. To selectively target GABAergic INTs and conditionally suppress gene expression of APP (and its homolog APLP2 to avoid potential compensatory mechanisms), we took advantage of a Cre-recombinase-based viral strategy in APP/APLP2 double floxed mice (Mehr et al., 2020). We examined the structural and functional consequences of deleting APP and APLP2 in CA3 GABAergic INTs at both cellular and circuit levels. We performed *ex vivo* electrophysiological recordings to explore whether subpopulations of GABAergic INTs are particularly vulnerable to APP/APLP2 deletion, and to examine changes in the nature of inhibitory inputs onto CA3 PCs. Additionally, using *in vivo* electrophysiological recordings in head-fixed awake mice, we highlighted disturbed oscillatory coupling between theta phase and low gamma amplitude following deletion of APP/APLP2 in GABAergic INTs. The theta-phase modulation of CA3 PC spiking and the strength of specific INT-to-INT connections were also reduced. Altogether, these findings highlight the importance of APP in CA3 GABAergic INTs in maintaining normal oscillatory processes and associated excitatory neuronal firing.

## Methods

### Experimental model

APP^fl/fl^/APLP2^fl/fl^ mice were obtained by crossing single APP^fl/f^ with APLP2^fl/fl^ homozygotic mouse lines originally obtained from Ulrike Müller’s lab (University of Heidelberg, Germany). These genes were flanked independently with 3 LoxP sites present between exon1 and the neoR cassette upstream of the start codon (Mallm et al., 2010). For the control condition, C57BL6/N mice were obtained from Charles River. Animals were bred and housed in the “Pole in Vivo” facility of the Interdisciplinary Institute for Neuroscience, following the regulations of the University of Bordeaux/CNRS Animal Care and Use Committee (authorization numbers #10946, #29169 and #37890). Both male and female mice were used in the course of this study, without taking in consideration the female’s estrous cycle. Throughout their lives, mice were housed with littermates, and food and water were provided ad libitum. Plexiglass cages were kept in a temperature-regulated room on a 12 hours dark/light cycle.

### Viral transduction

Adeno-associated viral vectors (AAV9.mDLX.CRE-EGFP) containing Cre recombinase and EGFP reporter gene under the mDLX promoter (Dimidschstein et al., 2016) were produced by the viral vector facility of Bordeaux “TransBioMed”. The titer of the viral construct ranged from 2.9 to 3.2E+13 particles/mL. To correctly adjust the volume and rate of infection, the virus was diluted to 1:5 into filtered PBS prior to stereotaxic injections. Mice aged approximately P25 were bilaterally injected under isoflurane anesthesia (1-4%) into the CA3 region (targeting CA3c/CA3b), with the following coordinates: AP -2.2, ML +/-2.5, DV -2.35. Those coordinates were established taking bregma as the reference point (zero offset). For *ex vivo* electrophysiological experiments, 400nL of viral solution was injected at a rate of 75nL/min using a precision micro pump (World Precision Instruments). For *in vivo* recordings on awake mice, 500nL of virus was unilaterally injected in order to slightly enlargen the site of infection. All animals were subcutaneously injected with buprenorphine and meloxicam to prevent post-surgery pain and carefully monitored for 2 days following surgery. Experiments on injected animals were made at least 3 weeks after surgeries, required time for proper level of expression of the transgene following neuronal viral infection.

### *Ex vivo* acute slice electrophysiology

On the morning of the recording day, mice were deeply anesthetized with ketamine (100mg/kg of body weight) and xylazine (10mg/kg) via intraperitoneal injection and then rapidly intracardially perfused with ice cold oxygenated (carbogen - 95% O2, 5% CO2) artificial cerebrospinal fluid (aCSF) containing: 120 mM NaCl, 16.5 mM glucose, 1 mM MgCl2 anhydrous, 2 mM CaCl2(H2O)2, 2.5 mM KCl, 1.25 mM NaH2PO4, 2.8 mM pyruvic acid, 27 mM NaHCO3, and 0.5 mM ascorbic acid. Osmolarity was set within a range of 305-315mOsm. Upon perfusion, brains were quickly dissected, and parasagittal slices (300 µm thick) were cut from both hemispheres using a vibratome (Leica Microsystems VT1200S, Germany). Subsequently, slices were kept for 20 minutes in a chamber of aCSF heated at 34°C. Slices were maintained oxygenated with carbogen at room temperature during the recording session (up to 6 hours).

Whole-cell patch clamp recordings were performed either from CA3 INTs (identified by shining blue light to visualize EGFP reporter emission in infected cells) or from CA3 PCs identified with a differential interference contrast microscope (Olympus BX51FI) with an immersion 60X objective (LUMPlanFI, N.A. 0.9). Borosilicate glass capillaries (Harvard apparatus) were pulled with a vertical pipette puller (Narishige) to a resistance of patch pipette of 3-6 MΩ. For current-clamp recordings, pipettes were filled with an internal solution containing the following: 115mM K-gluconate, 10mM KCl, 0.2mM EGTA, 0.02mM CaCl2, 10mM HEPES, 15mM phosphocreatine, 4mM MgATP, and 0.3mM NaGTP, with pH 7.3 using KOH and an osmolarity set to 300mOsm. For the majority of patch recordings made in voltage-clamp (VC), an intracellular solution with the following composition was used: 125mM Cs-methanesulfonate, 2mM MgCl2(H2O)6, 4mM NaCl, 10mM EGTA, 10mM HEPES, 5mM phosphocreatine, and 4mM NaATP, with pH 7.3 using CsOH and an osmolarity of 300mOsm. In order to record spontaneous (sIPSCs) and miniature inhibitory postsynaptic currents (mIPSCs) in the voltage-clamp mode at -70mV holding potential, a Cesium-chloride intracellular solution containing QX-314 Na^+^ channel blocker was specifically used, composed of: 130mM Cs-chloride, 100 mM NaCl, 10mM HEPES, 1mM EGTA, 0.1mM CaCl2, 10 mM phosphocreatine, 4 mM ATP-Mg, 0.4 mM GTP-Na, 5mM QX-314, with pH 7.2 using CsOH and an osmolarity of 285-290mOsm. Neurons were kept at a holding potential of -70mV, and leak current as well as series resistance were continuously monitored throughout each experiment. Signals were amplified using a HEKA EPC10 amplifier (HEKA Elektronik, Germany), filtered at 3.3kHz and digitized at 10kHz via PatchMaster Software (Germany). Recordings from cells with leak currents > 250pA and series resistance > 25MΩ were discarded. Moreover, neurons whose series resistance varied above 30% along the course of the recording were also rejected. All data was analyzed using IGOR PRO 6.3 software (Wavemetrics, USA) and the extension tool NeuroMatic (Rothman & Silver, 2018).

For current-clamp experiments, the resting membrane potential (RMP) was measured immediately upon break-in, to avoid the effects of intracellular solution dialysis. To study the membrane properties, cell excitability and firing behavior of patched INTs, we injected hyperpolarizing and then depolarizing currents into the cell in a stepwise manner with 10pA increments. Input resistance (Rin) was calculated by linearly fitting the change in voltage against the negative injected currents. For the recording of evoked Mf-CA3 excitatory post-synaptic currents (EPSCs) and INT-dependent feedforward inhibition – inhibitory post-synaptic currents (IPSCs) in CA3 PCs, electrical stimulations were performed using a second glass pipette filled with aCSF which was placed in the zone where MFs emerge from the DG granule cell layer. The baseline stimulation frequency was set at 0.1Hz and the stimulation amplitude was adjusted to obtain partial failure in MF-CA3 PCs EPSCs, in order to get minimal stimulation conditions (Marchal & Mulle, 2004). For the study of inhibition/excitation (I/E) ratio, CA3 PCs were voltage-clamped at +10mV, the reversal membrane potential for cations, allowing the detection of IPSCs. Spontaneous and miniature IPSCs were recorded in voltage-clamp in presence of NBQX (10µM) and D-APV (50µM) to block AMPA and NMDA receptors, respectively.

### Ex vivo electrophysiological data - statistical analysis

Statistics and graphical representations were made with GraphPad Prism 10.1.0. software (GraphPad Software, La Jolla, CA, USA). Values were first tested to check whether the data followed a normal distribution (D’Agostino & Pearson test) and outliers were removed by using the Grubbs’ test (alpha set to 0.05). Data was presented as bar graphs containing the individual data points and error bars (mean ± SEM). When comparing two populations, data that passed the normality test was compared using an unpaired t-test, while non-normally distributed data was compared using the Mann-Whitney test. When comparing non-normally distributed, paired data, a Wilcoxon rank test was used. When comparing cumulative frequencies, a Kolmogorov Smirnov test was used. When comparing data sets involving two independent factors (genotype and stimulus number, e.g.), statistical differences were tested using a two-way ANOVA followed by a Šídák’s multiple comparisons post-hoc test. Differences among groups were considered significant at p<0.05. No statistical method was used to predetermine the sample size. The experimenter was blinded to the genotype of the mice during the experiments and primary analysis.

### Immunohistochemistry

Mice were injected with a mix of ketamine and xylazine then intracardially perfused with 50 ml PBS, followed by PFA 4% until complete stiffness. After an overnight fixation in PFA 4%, the brains were transferred to PBS. 50μm coronal slices were acquired with a vibratome (Leica, VT1200S, Germany). Sections were permeabilized and blocked in a 5% normal goat serum and 0.3% Triton X-100 solution for 2 hours at room temperature and then incubated in primary antibodies (APP C-terminal Y188 rabbit antibody Abcam #32136, Parvalbumin polyclonal guinea pig antibody Synaptic System #195004, Somatostatin monoclonal rat antibody Abcam #30788) overnight at 4°C under gentle agitation. After PBS washing (3 times), the slices were incubated with secondary antibodies for 2 hours at room temperature. Finally, the slices were mounted with Dapi-containing mounting solution (Vectashield, H-1200-10) and imaged with a confocal microscope (Leica DM6 CFS TCS SP8).

### Morphological reconstruction of CA3 INTs

After patch-clamp recording of CA3 INTs and intracellular infusion of biocytin at 0.2% (Sigma, B4261) through the patch pipette for roughly 15 minutes, the position of the recorded cells was annotated and the slices were transferred from the recording chamber to a PFA 4% filled plate. From each acute brain slice, a maximum of two CA3 INTs were recorded. After overnight fixation, free-floating slices were washed 3 times in PBS and then permeabilized and blocked at room temperature (RT) for 1 hour in a 5% normal goat serum and 0.3% Triton X-100 solution of PBS. Sections were incubated a Streptavidin-555 (Thermofisher S32355) solution (dilution 1/2000 in PBST) overnight at room temperature under agitation. Finally, brain sections were washed 1 time in PBST and 2 times in PBS and mounted with Vectashield mounting medium. The fluorescent signal of mounted brain slices was acquired with a confocal microscope (TCS SP5 Leica, immersion 20X objective). Neuronal reconstruction of the recorded CA3 INTs was achieved using the SNT (Simple Neurite Tracer) plugin in the Fiji/ImageJ software (National Institutes of Health, New York, USA). A Z-stack maximal projection was performed to follow the dendrites in x, y and z axes as well. The center of the soma was used as the point of reference from which the tracing of the dendrites was manually performed by following the fluorescent signal of each branch until reaching a signal not resolutive enough for the eye. After morphological reconstruction we performed a Sholl analysis. The description of the parameters used to study the dendritic morphology (e.g. complexity index (DCI), cable length, No. of branches) of the reconstructed cells is available online: https://imagej.net/plugins/snt/metrics.

### Clustering of recorded CA3 INTs

A custom script was used to perform the classification of the recorded CA3 INTs. It was written in R programming language and run in the RStudio integrated developing environment. The clustering was based on a t-SNE (t-distributed stochastic neighbor embedding) analysis with perplexity = 8 and number of iterations = 1000. The electrophysiological parameters used were the input resistance, the rheobase, the mean firing rate, the frequency-current gain and the SAG ratio.

### *In vivo* silicon probe electrophysiology

At around P30, APP^fl/fl^/APLP2^fl/fl^ or control C57BL6/N mice were unilaterally injected with the AAV9.mDLX-CRE-EGFP viral construct in the CA3 region and implanted with a head bar to allow for later head fixation. The head bar was fixed to the skull with dental cement. To facilitate the subsequent craniotomy, the region of the skull above the hippocampus was kept free from dental cement by a plastic ring attached with super glue and filled with silicone (Kwik-Sil, WPI). Sessions of habituation to the recoding setup were started after 5 to 7 days of post-implantation recovery. During the first session lasting 10 minutes, mice were not head fixed but freely moving within an enclosure on the running wheel to avoid high stress. Then, mice were progressively trained to remain head-fixed on the wheel by increasing the time of the habituation: from 10 minutes on the first session to 20, 40 and up to 1 hour after 4 days. The mice were then trained daily for 1 hour during at least 3 weeks counting from the surgery/implantation day. This period of time was selected to allow proper viral expression and to optimize the training period on the wheel. On the day of the recording, mice were anesthetized and the silicone was removed from the plastic ring to have access to the skull. A craniotomy using a drill was performed following an elliptic shape to fit the geometry of the 4-shank silicon probe and the bregma was used as point of reference. The craniotomy orientation along antero-posterior and medio-lateral axes was given by two marks: (1) -1.35 (AP); -2.00 (ML) (AP), (2) -3.05 (AP); -3.00 (ML). The brain surface was constantly maintained humid with a saline solution. Mice were allowed to recover for an hour and were then head-fixed for the recording session. A 4-shank silicon probe was inserted in CA3 (A4x16-poly2-5mm-23s-200-177) connected to a 64-channel adaptor (Neuronexus, Adpt-A64-OM32x2-sm). The adaptors were connected to Intan headstages (Intan RD 64) and then connected to an Intan 512-ch RHD Recording Amplifier. The wheel systems were provided by Polywheel (Imetronic, France). Before starting the recording, the probe was dipped in a DiI-based solution in order to mark the probe path and the recording site. The probe was inserted in the brain at a speed not exceeding 1μm/second using a micromanipulator (Scientifica). The assessment of the location was performed by constant monitoring of depth and of the brain signal. When electrophysiological signals were consistent with a localization in CA3, the recording session was started and lasted 30 minutes. At the end, the probes were taken out and the mice were euthanized for post-hoc brain histology.

### Histology (post *in vivo* silicon recording)

Following in vivo recordings, mice were deeply anesthetized with isoflurane, dislocated and their brain was dissected in order to evaluate the level of viral infection within CA3 region and to confirm the correct probe location at the recording site in the brain. Following the extraction, the mouse brain was submerged in PFA 4% solution for approximately 24 hours. Then, 50μm coronal slices were made with a vibratome (Leica, VT1200S, Germany), slices were mounted with Dapi-containing mounting solution (Vectashield, H-1200-10) and imaged with regular epifluorescence microscope (Nikon Eclipse Ni, 10X dry objective N.A. 0.30).

### *In vivo* electrophysiological data - quantification and statistical analysis

#### Local field potentials

Raw data files (.dat format) recorded at 20kHz for LFP were read into Python using the Neo package (Garcia et al., 2014) and signal processing was performed with custom routines using the SciPy package (Virtanen et al., 2020). LFP signal was down-sampled 16 times to 1250 Hz and filtered using a 4-pole Butterworth band-pass filter between 0.2 Hz and 300 Hz.

#### Spike sorting

***S***pike sorting was performed in Kilosort4 (Pachitariu et al., 2024) and hand-curated in Phy (https://github.com/kwikteam/phy). Unit classification into putative pyramidal cells and interneurons was based on waveform shape and firing characteristics (Csicsvari et al., 1999). Several measures were calculated and those that provided the best separation between two groups were used. Pyramidal cells had a wide waveform (trough-to-peak of at least 0.5ms) and a low average firing rate (less than 8 Hz). Interneurons had a narrow waveform (trough-to-peak of less than 0.8ms) and a low propensity to burst (less than 30% of interspike intervals <6ms). The last useful criterion was the width of the autocorrelation, another measure of the cell’s propensity to fire at close intervals, and defined as the lag when the autocorrelation fell below its mean. Interneurons had autocorrelation widths greater than 24ms. With these criteria, a portion of the population didn’t fall into either category and was excluded from some analyses. Subsequent analysis of putative downstream interactions confirmed that units labeled as pyramidal has mostly excitatory downstream interactions, and interneurons had mostly inhibitory.

#### Brain state classification

Detection of theta epochs was performed on one LFP trace taken from a channel manually determined to be in the stratum radiatum of CA3. Individual theta cycles were detected following the procedure found in a published paper by Vitor Lopes-dos-Santos and colleagues (Lopes-dos-Santos et al., 2018), using an empirical mode decomposition (EMD) filter and finding the individual oscillation cycles that crossed an amplitude threshold (median of the absolute value of the filtered theta trace) and matched theta frequency (4-12 Hz). Consecutive theta cycles occurring within a 250ms time window were joined into theta epochs and only theta epochs longer than 550ms were included in subsequent analyses.

#### Population activity

Two measures of population activity in CA3 were used: pairwise correlation coefficients and population activity events. First, spike trains were obtained by binning the firing of CA3 PCs into 20ms bins, allowing to calculate the correlation coefficient for each pair of neurons. Higher pairwise correlation coefficients indicated that the pair of cells tended to fire within the same 20ms time bins. Spike trains of CA3 PCs were also analyzed to study the population events during recordings. From the 20ms spike binning, the number of PCs concomitantly active was counted. Then we calculated a population event threshold (in number of active cells) by session. For each single session, the spike trains were shuffled and the 95% confidence intervals around the distribution of the number cells simultaneously active over time bins was calculated. We did the spike train shuffling as follows: for each shuffle, all the spike trains of the simultaneously recorded cells were shifted by a different random number of time bins, with activity of the latter part of the train being wrapped around to the beginning. This ensured that individual spike trains were shifted with respect to each other while still keep the original spiking patterns of each individual cell. The number of shuffling iterations was set to 100 times, producing the 95% confidence interval around the number of simultaneously active cells. For each recording, the threshold was defined as the smallest number of simultaneously active cells that occurred more often in the real dataset than the 95% confidence interval of the shuffled dataset. Time bins that had more active cells than the threshold was counted as a population event, as more cells were active than were expected by chance. Lastly, we calculated the proportion of “burst-like” population events. The maximal inter-event interval (iei) was set to 200ms (gathering 10 time-bins) to consider consecutive population events having a bursty behavior with temporal firing synchronicity.

#### Theta spike modulation

To study the modulation of firing due to theta oscillations, we first filtered the LFP from one channel manually determined to be in the stratum pyramidal of the CA3 using a 4-pole butterworth filter with bandpass cutoff frequencies of 4 and 12 Hz. The instantaneous phase of theta at the time of each spike was determined using the Hilbert transform. Only spikes occurring during the theta epochs determined as described above were used in this analysis. For each cell unit, the resultant vector length (circ_r) and the mean direction (circ_mean) were extracted from the distribution of spike theta phases by using the pingouin python package (https://pingouin-stats.org/). A higher mean vector length indicates a less uniform spike distribution across theta phases and is interpreted as a stronger theta modulation. The mean vector length and the preferred phase were used to assess the level of theta spike modulation and whether it was changed between conditions.

#### Cross frequency coupling

LFP traces from the CA3 stratum radiatum were used to analyze theta and gamma spectral components of the LFP signal. For each recording session, the phase-amplitude coupling (PAC) was estimated by computing theta-gamma comodulograms with the Pactools python package (Dupré la Tour et al., 2017) (https://pactools.github.io/index.html#) applying the ‘tort’ method (Tort et al., 2010). The low frequency range (from 1 to 16Hz) was binned into 0.2Hz frequency bins and the high frequency range (from 16 to 160Hz) into 2Hz frequency bins. The modulation index (MI) values obtained for each theta cycle were normalized to the max MI from one recording/animal. Averaged comodulograms within theta frequency range restricted to 5-8Hz were used to find the theta modulation peaks for both low (16-40 Hz) and high (40-110 Hz) gamma.

#### Cell to cell connectivity analysis

To evaluate the spiking influence of an upstream ‘presynaptic’ cell onto a downstream ‘postsynaptic’ one (cell to cell connectivity) among putative CA3 PCs and INTs, we analyzed the temporal relationship of spikes between cell units. Only units with an average firing rate of at least 0.2 Hz were used for this analysis, as insufficient firing led to unreliable cross-correlograms. We used a method similar to that described in (Senzai & Buzsáki, 2017). For each cell pair, a cross-correlogram (CCG) (1ms time bins from -50 to 50ms) was computed by summing the number of spikes of the postsynaptic partner found within each lagtime bin with respect to the firing of the presynaptic partner. The pair was considered to be functionally connected when the CCG at short lag times was significantly higher or lower than what would be expected by chance, for excitatory and inhibitory interactions, respectively. The critical window to study ‘monosynaptic’ cell connectivity was set to 1-5ms for excitatory connections and to 1.5-5ms for inhibitory ones After having applied a gaussian filter (sigma = 10ms) on the cross-correlogram, the Poisson distribution was used to define the maximum and minimum values expected by chance (at 0.999999 and 0.00001 in the cumulative Poisson distribution). The strength of the connection was assessed by calculating the area under the curve (AUC) within the critical window between the CCG and the gaussian-filtered CCG, normalized by the area between the threshold and the gaussian-filtered CCG. The proportion of connection for each type of connection (i.e. PC onto INT, PC onto PC, INT onto PC and INT onto INT) was calculated by dividing the number of experimentally found connectivity by the theoretical maximal number of connections given the number of identified cell units of each type in the session.

#### Statistical analysis

Parametric and nonparametric statistics were performed with the SciPy package (Virtanen et al., 2020) and circular statistics with the pycircstat2 package (https://github.com/circstat/pycircstat2). Unless otherwise noted, data were expressed as the median +/-the mean absolute derivation (MAD) about the median, which was calculated by subtracting each data point from the group’s median, taking the absolute value, summating, and then dividing by the number of data points. These measures of central tendency and variation were used because many distributions were found not to be normal. A Mann-Whitney U test was performed to compare independent groups (control versus APP/APLP2dKO CA3 INTs). To compare two non-independent groups, as in comparing the non-theta and theta states within one animal, a Wilcoxon signed-rank test was used. In the case of multiple comparisons, the Benjamini-Hochberg correction was used to reduce the occurrence of type II errors (false negative results). For all figures, significance of p<0.05 was represented with *, p<0.01 with **, p<0.001 with ***, and p<0.0001 with ****.

## Results

### Morpho-functional properties of subpopulations of CA3 INTs lacking APP/APLP2

In order to study the physiological role of APP in INTs, we performed a targeted genetic deletion of APP and of its homologous APLP2 proteins to avoid compensatory mechanisms (Fanutza et al., 2015; Mehr et al., 2020) by stereotaxic injection of the AAV9.mDLX-CRE-EGFP viral construct in the dorsal CA3 region of APP^fl/fl^/APLP2^fl/fl^ mice (referred to APP/APLP2 cKO mice). The use of the mdlx promoter allowed the expression of the Cre-recombinase fused to the enhanced-green fluorescent protein (used as a marker of infection) selectively in GABAergic INTs (Dimidschstein et al., 2016) (Fig. 1A and B). For control conditions, the same construct was injected in wild type (C57B6/N) mice. We confirmed that CA3 INTs express APP using immunostainings on hippocampal fixed brain slices in control mice (non-injected APP^fl/fl^/APLP2^fl/fl^ mice). We calculated the proportion of CA3 GABAergic INTs among two dominant INT populations (i.e. parvalbumin (PV) and somatostatin (SST) expressing neurons) which were immunopositive for APP (Fig. S1A and B). In the dorsal CA3 region, we found that approximately 84% of PV-positive neurons colabelled for APP, and 100% of SST-positive neurons colabelled for (Fig. S1C). The AAV9.mDLX-CRE-EGFP viral construct through the mdlx promoter targets all types of GABAergic INTs (Dimidschstein et al., 2016), as illustrated for PV+ INTs (Fig. S1D).

**Figure 1.**
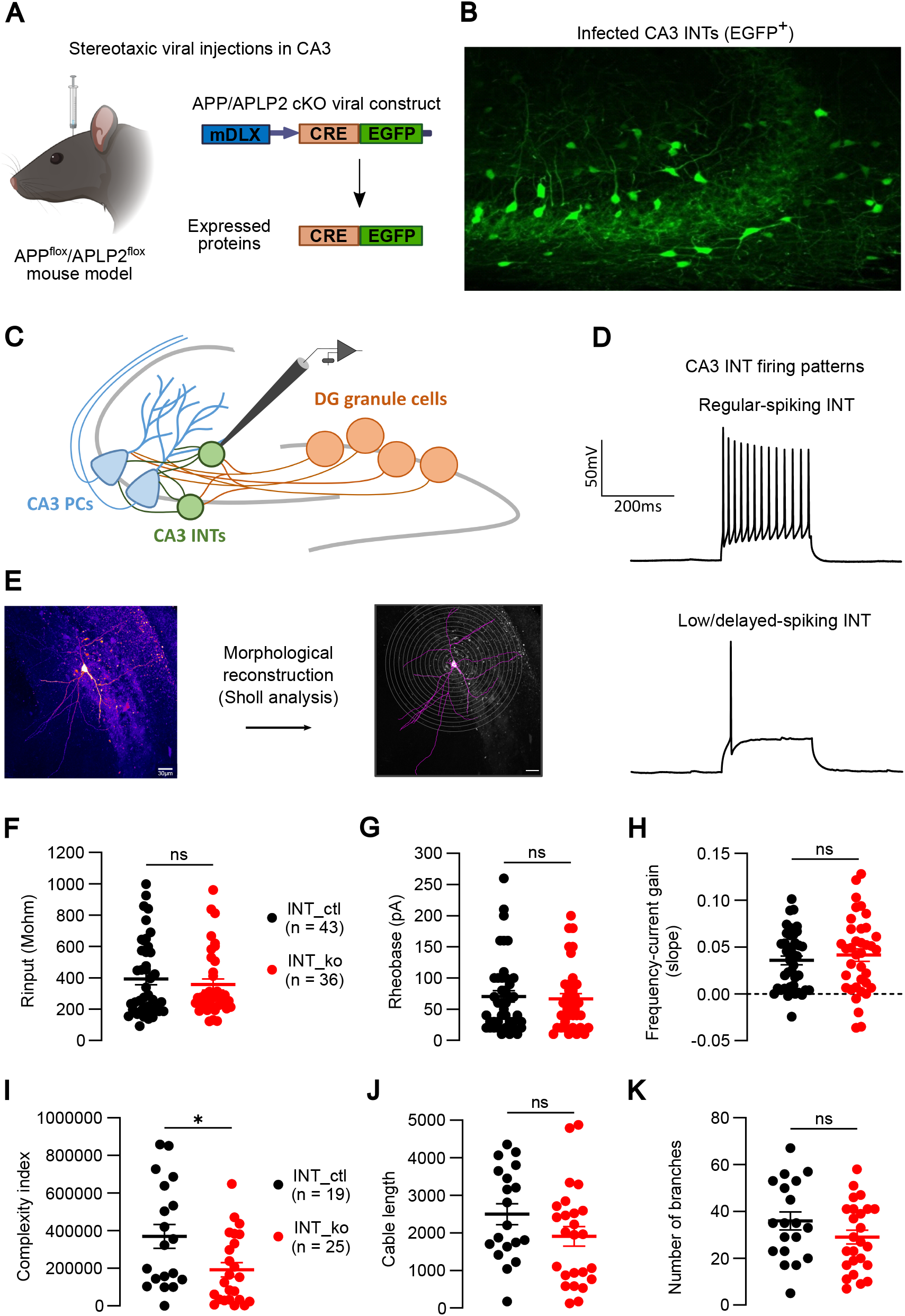
The selective deletion of APP/APLP2 in CA3 INTs only has minor effects on intrinsic properties. **A.** Viral strategy using an AAV9.mDLX-CRE-EGFP viral construct in order to selectively express Cre-recombinase under the control of the mdlx promoter, leading to deletion of both APP and APLP2 induced in GABAergic INTs. Bilateral viral injections were performed in the CA3 hippocampal region of APP^fl/fl^/APLP2^fl/fl^ mice (experimental group). Wild type (C57B6/N) mice were used for control. **B.** Confocal image of infected CA3 INTs expressing EGFP in hippocampal slices of APP/APLP2 cKO mice. **C.** Current-clamp recordings were performed in control (only EGFP-expressing INTs) and in APP/APLP2 cKO (and EGFP-expressing) CA3 INTs. **D.** Example traces of AP discharge in a typical regular-spiking INT (top) and of a low/delayed-spiking INT (bottom) in response to a 250ms depolarizing current step. **E.** Morphological reconstruction of patched CA3 INTs. Left: confocal image of a biocytin-filled CA3 INT. Right: Sholl analysis of reconstructed dendritic arborization. Neuronal electrophysiological features (i.e. membrane properties and cell excitability) of CA3 INTs, such as input resistance (**F**), rheobase (**G**) and frequency-current (F/I) gain (**H**) are not changed between conditions (n=43 for INT_ctl; n=36 for INT_ko; Mann-Whitney test: ns = not significant). The input resistance was quantified by linearly fitting the level voltage fluctuation against the injected current. The rheobase corresponds to the minimal amount of current required to elicit APs. The F/I gain slope was obtained by linear regression of the number of APs evoked by increment of current steps, from hyperpolarizing to depolarizing. There is a significant reduction of the complexity index (**I**) in INT_ko compared to INT_ctl (369459 ± 63774 for INT_ctl ; 192429 ± 37543 for INT_ko ; unpaired T test: p = 0.0163), which is not the case for other morphological parameters such as cable length (**J**) and number of branches (**K**). Data are represented as mean ± SEM, with individual dots matching each recorded CA3 INT.

We first investigated the morphological and functional properties of INTs lacking APP/APLP2. CA3 INTs were recorded in the current-clamp mode with an intracellular solution containing biocytin (2mg/mL) for post-recording neuronal visualization and reconstruction (Fig. 1C and E). Single action potentials (APs) and trains of APs were recorded by injecting either short 5ms positive current pulse or 200ms hyperpolarizing and depolarizing currents in a step-wise manner (from -80 to 210pA). Upon positive current steps, distinct firing patterns were recorded among the patched INTs (Fig. 1D). The membrane parameters (e.g. input resistance) and properties of excitability (i.e. rheobase, frequency-current (F-I) gain) were not changed between genotypes (Fig. 1F-H). After a biocytin-streptavidin based neuronal revelation, slice fixation and confocal image acquisition of the recorded CA3 INTs, we performed a Sholl analysis to study neuronal morphology (Fig. 1E). The group comparison of the Sholl parameters only showed a significant difference in the complexity index of the dendritic arborization (Fig. 1I-K, mean value: 369459 ± 63774 for INT_ctl (n=19); 192429 ± 37543 for INT_ko (n=24); unpaired T test: p=0.0163), pointing to a reduced dendritic coverage in APP/APLP2 cKO mice as compared to control.

The hippocampal GABAergic INTs are composed of various cell subtypes forming a heterogeneous population with distinct morphology, electrophysiological properties, spatial distribution and neuronal compartment targets (Tzilivaki et al., 2023). This diversity was experimentally sensed by the heterogeneity of the recorded morpho-functional properties in our dataset. To test whether certain subpopulations of INTs are particularly vulnerable to the deletion of APP/APLP2, we performed a dimensionality reduction analysis based on the quantified electrophysiological parameters (i.e. input resistance, rheobase, F-I gain, peak firing ratio and SAG ratio). The t-distributed stochastic neighbor embedding (t-SNE) analysis, that associates the close proximity in the 2D plot vector distribution to a similarity of neuronal features, allowed identifying three main clusters of CA3 INTs, here designted as cluster A, cluster B and cluster C (Fig. 2A). The analysis of the cell location on hippocampal slices (CA3 layer of the patched INTs) did not show any dominant layer among the three clusters (Fig. 2B, right). The inter-cluster comparison of the electrophysiological parameters highlighted as expected significant differences between the clusters A, B and C (Fig. 2B-D, mean input resistance: 192.7 ± 10.7MΩ for cluster A (n=21), 285.8 ± 17.0MΩ for cluster B (n=35), 685.0 ± 33.3MΩ for cluster C (n=23), Kruskal-Wallis multiple comparison: p=0.0197 cluster A *vs* cluster B, p<0.0001 cluster A *vs* cluster C, p<0.0001 cluster B *vs* cluster C; mean rheobase: 158.1 ± 15.0pA for cluster A, 54.9 ± 4.9pA for cluster B, 28.8 ± 3.3pA for cluster C, Kruskal-Wallis multiple comparison: p<0.0001 cluster A *vs* cluster B, p<0.0001 cluster A *vs* cluster C, p=0.0188 cluster B *vs* cluster C; mean F-I gain: 0.0076 ± 0.0015 for cluster A, 0.0654 ± 0.0046 for cluster B, 0.0259 ± 0.0066 for cluster C, One-way ANOVA multiple comparison: p<0.0001 cluster A *vs* cluster B, p=0.0179 cluster A *vs* cluster C, p<0.0001 cluster B *vs* cluster C). These latter results validate the identification of three different subpopulations of recorded INTs with distinct passive and active membrane properties, which were not associated with morphological changes (Fig. S2A-C, no difference in cable length, number of branches and complexity index between the three clusters of cells). To study the possible impact of APP/APLP2 conditional deletion on each cluster, we ran an intra-cluster group comparison. The input resistance was significantly increased for INT_ko compared to INT_ctl in cluster A (Fig. 2E, mean input resistance: 175.0 ± 11.98MΩ for INT_ctl (n=13); 221.4 ± 16.39MΩ for INT_ko (n=8); Mann-Whitney test: p = 0.0302 INT_ctl *vs* INT_ko), whereas the other electrophysiological properties were not modified (Fig. 2F and G). Although the low number of cells in clusters A and B limited the statistical power to detect significant differences in morphological parameters, our data suggest a subtle but consistent reduction of the dendritic length and complexity in INT_ko compared to INT_ctl in cluster A (Fig. S2D-F, mean cable length: 2754 ± 527µm for INT_ctl (n=5), 1274 ± 351µm for INT_ko (n=3); mean number of branches: 46.4 ± 7.8 for INT_ctl, 18.0 ± 1.0 for INT_ko; mean complexity index: 526747 ± 133386 for INT_ctl, 59197 ± 14691 for INT_ko). No significant difference was observed in cluster C between control and cKO INTs (Fig. S2D-F, mean cable length: 2010 ± 433µm for INT_ctl (n=9), 1605 ± 335µm for INT_ko (n=8); mean number of branches: 35.8 ± 5.5 for INT_ctl, 25.8 ± 4.1 for INT_ko; mean complexity index: 260609 ± 80923 for INT_ctl, 167409 ± 60464 for INT_ko, Mann-Whitney test: p = 0.4807).

**Figure 2.**
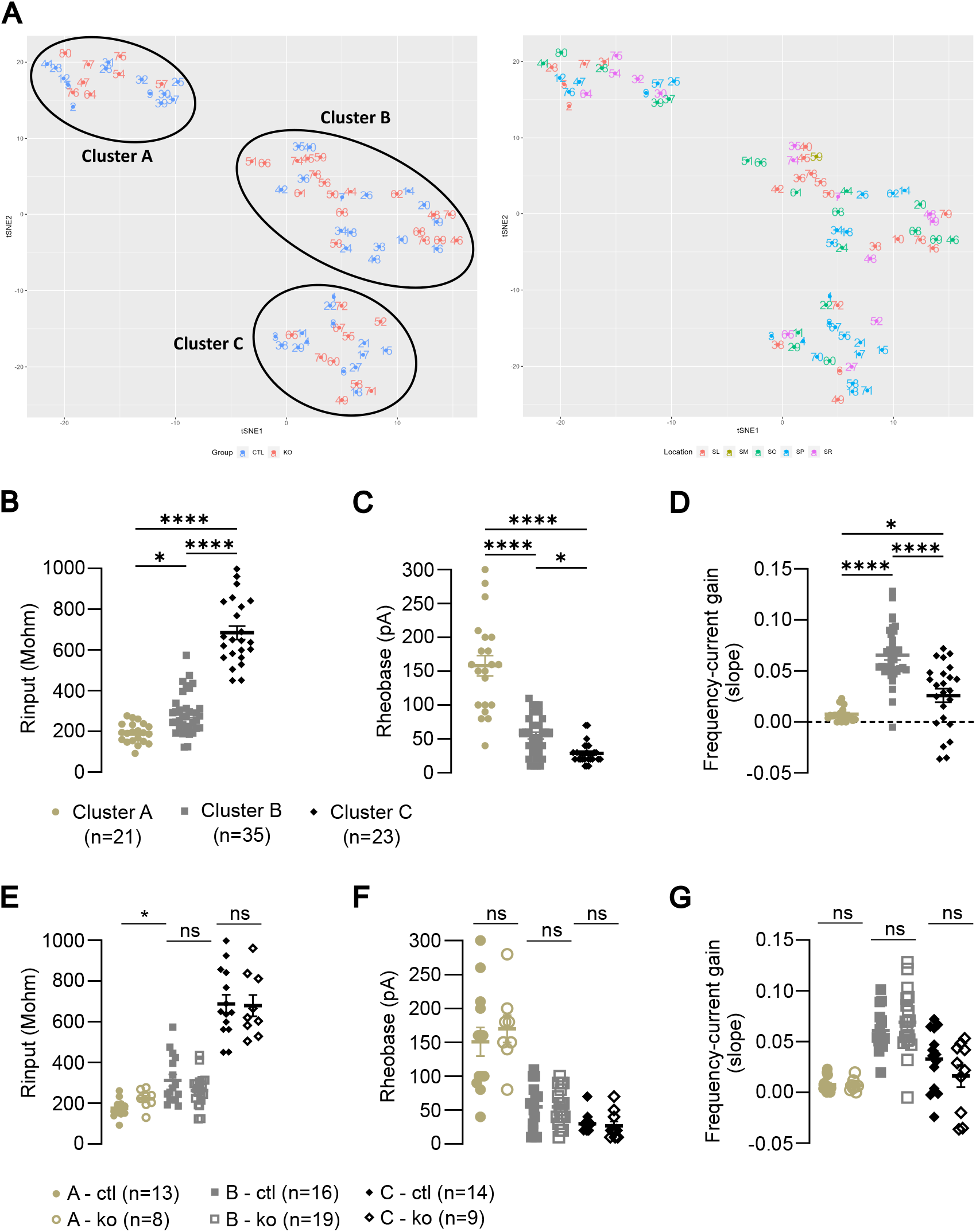
Characterization of morphofunctionally distinct subpopulations of CA3 INTs in control and cKO conditions. A. A multidimensional reduction analysis (t-SNE) was applied to the dataset. Left: t-SNE analysis 2D output with identification of three main clusters of INTs based on electrophysiological parameters. Clusters on the plot include both control (blue dots) and APP/APLP cKO (red dots) CA3 INTs. Right: Location in CA3 area of the recorded cells among the clusters, sorted by color code: stratum lucidum (SL - red), stratum moleculare (SM – olive green), stratum oriens (SO - green), stratum pyramidale (SP - blue) and stratum radiatum (SR - pink), respectively. Significant differences are observed between clusters of cells, confirming the identification of INT subpopulations with distinct neuronal properties, namely input resistance (B), rheobase (C) and F-I gain (D). Data are represented as mean ± SEM, with individual dots matching each recorded CA3 INT. Statistics: mean input resistance: 192.7 ± 10.7MΩ for cluster A (n=21), 285.8 ± 17.0MΩ for cluster B (n=35), 685.0 ± 33.3MΩ for cluster C (n=23), Kruskal-Wallis multiple comparison: p=0.0197 cluster A vs cluster B, p<0.0001 cluster A vs cluster C, p<0.0001 cluster B vs cluster C; mean rheobase: 158.1 ± 15.0pA for cluster A, 54.9 ± 4.9pA for cluster B, 28.8 ± 3.3pA for cluster C, Kruskal-Wallis multiple comparison: p<0.0001 cluster A vs cluster B, p<0.0001 cluster A vs cluster C, p=0.0188 cluster B vs cluster C; mean slope F-I gain: 0.0076 ± 0.0015 for cluster A, 0.0654 ± 0.0046 for cluster B, 0.0259 ± 0.0066 for cluster C, One-way ANOVA multiple comparison: p<0.0001 cluster A vs cluster B, p=0.0179 cluster A vs cluster C, p<0.0001 cluster B vs cluster C. The specific deletion of APP/APLP2 impacted the input resistance (E) of INTs only in cluster A (mean Rinput: 175.0 ± 11.98MΩ for INT_ctl (n=13); 221.4 ± 16.39MΩ for INT_ko (n=8); Mann-Whitney test: p = 0.0302 INT_ctl vs INT_ko, ns = not significant), contrary to clusters B and C. No differences were observed for the rheobase (F) and F-I gain (G) among the three clusters. Statistics: cluster B: n=16 INT_ctl, n=19 INT_ko; cluster C: n=14 INT_ctl, n=10 INT_ko; Mann-Whitney or unpaired T test: ns = not significant.

This set of experiments, due to a lack of neurochemical characterization (unexploitable results obtained from RNA-sequencing) coupled with divergent morpho-functional properties of the recorded CA3 INTs, as exemplified by the dominant firing patterns and fluctuating cell morphology found in cluster A (Fig. S3), did not allow us to precisely determine the cell identity (classical INT subtypes) of the three clusters. Altogether, our results indicate that the deletion of APP/APLP2 from GABAergic INTs from CA3, only has minor consequences on intrinsic morphofunctional properties.

### Deletion of APP/APLP2 from CA3 GABAergic INTs modifies the properties of spontaneous inhibitory events onto CA3 PCs

To test the potential functional impact of APP/APLP2 cKO on the synaptic coverage of CA3 PCs, we recorded both spontaneous and miniature (in presence of TTX 1µM) inhibitory post-synaptic currents (sIPSCs and mIPSCs, respectively) in CA3 PCs (Fig. 3A). We recorded IPSCs in the presence of QX314 (a voltage-dependent sodium channel blocker) in the intracellular solution, and NBQX and D-APV (antagonists of AMPA and NMDA receptors) in the aCSF (Fig. 3B). The mean frequency of both sIPSCs and mIPSCs were not significantly different between genotypes (Fig. 3C and G). However, albeit no change in the average mean amplitude was found (Fig. 3D and H), we observed a significant change in the cumulative distribution of the amplitudes of sIPSCs and mIPSCs, with a shift towards bigger values in APP/APLP2 cKO mice (INT_ko) compared to control CA3 INTs (INT_ctl) (Fig. 3D and H, mean amplitude of spontaneous events: 38.3 ±0.5pA for INT_ctl (n=18), 46.9 ± 0.6pA for INT_ko (n=17), Kolmogorov-Smirnov test: p < 0.0001; mean amplitude of miniature events: 32.1 ± 0.1pA for INT_ctl (n=17), 37.3 ± 0.3pA for INT_ko (n=15), Kolmogorov-Smirnov test: p < 0.0001). This was associated with a significant decrease of the rise time of sIPSCs and mIPSCs in APP/APLP2 cKO mice (Fig. 3E and I, mean rise time sIPSCs: 3.0 ± 0.1ms for INT_ctl (n=18), 2.4 ± 0.2ms for INT_ko (n=16), Mann-Whitney test: p = 0.0086; mean rise time mIPSCs: 2.7 ± 0.1ms for INT_ctl (n=17), 2.1 ± 0.1ms for INT_ko (n=14), unpaired T test: p = 0.0011), with cumulative distributions shifted towards shorter rise times for both spontaneous and miniature inhibitory events (Fig. 3E and I, mean rise time of spontaneous events: 2.7 ± 0.03ms for INT_ctl, 2.3 ± 0.03ms for INT_ko, Kolmogorov-Smirnov test: p < 0.0001; mean rise time of miniature events: 2.5 ± 0.02ms for INT_ctl, 2.1 ± 0.02ms for INT_ko, Kolmogorov-Smirnov test: p < 0.0001). Interestingly, by plotting the mean amplitude of events against the mean rise time for each recorded CA3 PC, we observed a modified plot distribution with a trend towards an increased amplitude associated with a reduced rise time of sIPSCs within cells for the INT_ko but not for INT_ctl (Fig. 3F). Perisomatic inhibition on principal neurons generates postsynaptic currents with larger amplitude and faster rise time to the soma (i.e. patch-pipette site of recording), while inhibitory contacts onto distal dendrites elicit currents that undergo attenuation along the dendritic route, with reduced amplitude and longer rise time when integrated at the soma (Kubota et al., 2015). We thus suspect that changes in the kinetics and amplitude of spontaneous IPSCs are caused by a circuit reorganization of CA3 GABAergic axon terminals, with an increase in connections targeting preferentially the somato-dendritic compartment of CA3 PCs.

**Figure 3.**
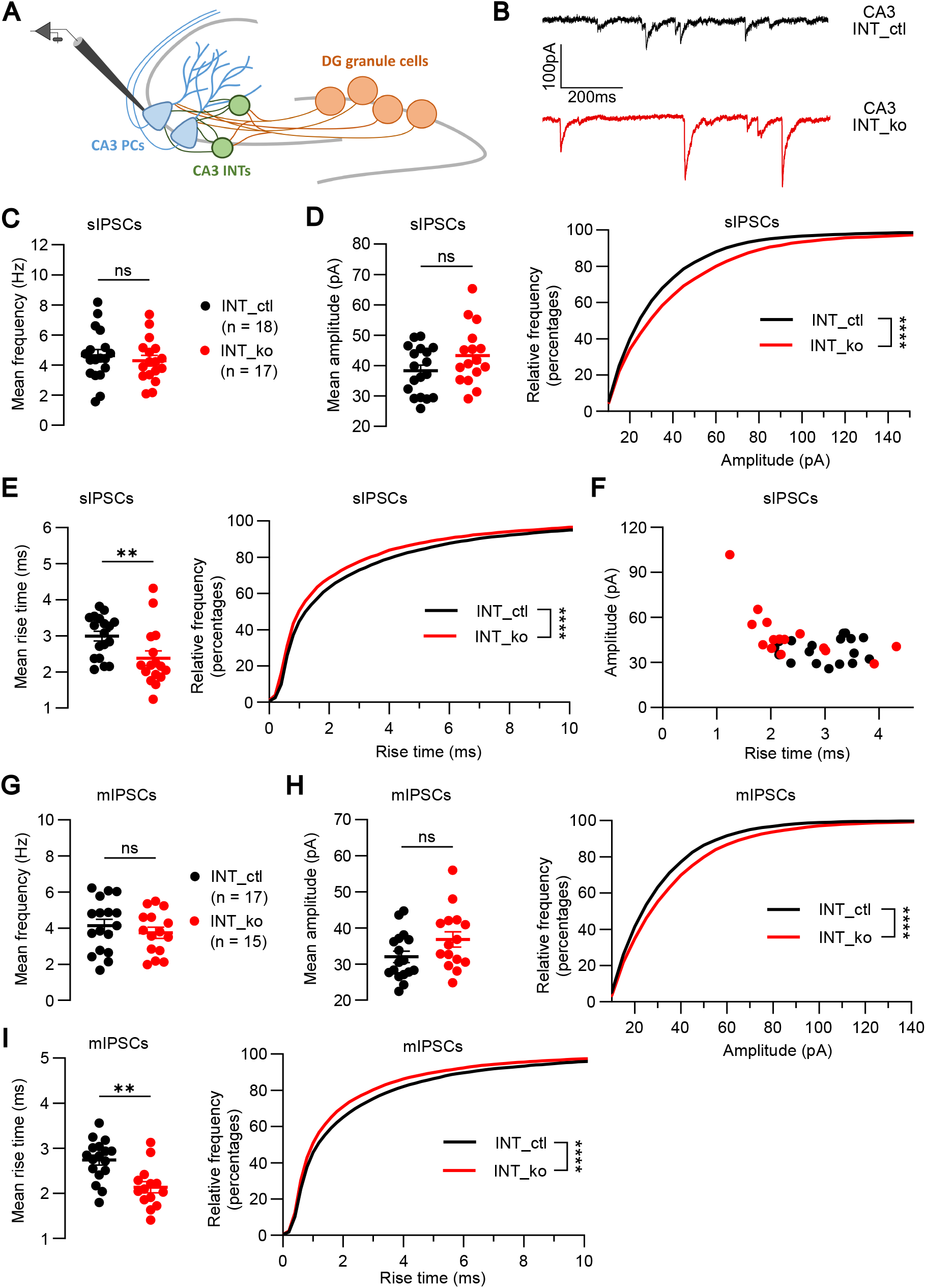
The selective and conditional deletion of APP/APLP2 in CA3 INTs modifies the properties of spontaneous inhibitory events onto CA3 PCs. A. Patch-clamp recordings of IPSCs and mIPSCs in CA3 PCs. The holding potential was set at -70mV and GABA-mediated inhibitory events were isolated using a cesium-chloride based intracellular solution containing QX-314 (5mM). NBQX (10µM) and D-APV (50µM) was present in the ACSF in order to antagonize AMPA, kainate and NMDA receptors. B. Example traces of recorded sIPSCs in control (top) and APP/APLP2 cKO (bottom) conditions. C and D. No significant difference is observed in the mean frequency and amplitude of sIPSCs between genotypes (INT_ctl: n = 18; INT_ko: n = 17; statistics: unpaired T test, ns = not significant). However, a shift in the cumulative distribution of amplitudes of spontaneous events is observed in INT_ko as compared to INT_ctl (mean amplitude sIPSCs: 38.3 ± 0.5pA for INT_ctl, 46.9 ± 0.6pA for INT_ko, Kolmogorov-Smirnov test: p < 0.0001; bin width = 5). E. A decreased rise time of sIPSCs in cKO mice (mean rise time sIPSCs: 3.0 ± 0.1ms for INT_ctl, 2.4 ± 0.2ms for INT_ko, Mann-Whitney test: p = 0.0086) is associated with a significant shift in the cumulative distribution of sIPSCs towards shorter rise times (mean rise time: 2.7 ± 0.03ms for INT_ctl, 2.3 ± 0.03ms for INT_ko, Kolmogorov-Smirnov test: p < 0.0001, bin width = 0.2). F. Plotting of the mean amplitude and rise time of the recorded cells. G and H. The mean frequency and amplitude of mIPSCs recorded in presence of TTX (1µM) are not significantly different between genotypes (INT_ctl: n = 17; INT_ko: n = 15; statistics: unpaired T test, ns = not significant). The cumulative distribution of mIPSCs is significantly shifted towards higher amplitudes in INT_ko as compared to INT_ctl (mean amplitude: 32.1 ± 0.1pA for INT_ctl, 37.3 ± 0.2pA for INT_ko, Kolmogorov-Smirnov test: p < 0.0001; bin width = 5). I. This is associated with a decreased rise time of mIPSCs in cKO mice (mean rise time: 2.7 ± 0.1ms for INT_ctl, 2.1 ± 0.1ms for INT_ko, unpaired T test: p = 0.0011) and with a significant change in their cumulative distribution (mean rise time: 2.5 ± 0.02ms for INT_ctl, 2.1 ± 0.02ms for INT_ko, Kolmogorov-Smirnov test: p < 0.0001; bin width = 0.2).

### Feedforward inhibition and inhibition/excitation in CA3 PCs in absence of APP/APLP2 in INTs

We next assessed whether modifications in GABAergic transmission in APP/APLP2 cKO mice (indirectly observed during spontaneous activity) had an impact on feedforward inhibition (FFI) evoked by the stimulation of mossy fibers (MFs) originating from the DG cells, and consequently influenced the overall excitation/inhibition balance at a cellular level. MFs form direct excitatory inputs onto CA3 PCs but also contribute to indirect, disynaptic feedforward inhibition of CA3 PCs by contacting local interneurons through filopodia (Lawrence & McBain, 2003; Rebola et al., 2017). We recorded monosynaptic MF-CA3 EPSCs or disynaptic CA3 INT–PC IPSCs evoked by the stimulation of the MF track with a stimulation electrode positioned in the molecular layer of the DG. At basal transmission, MF-CA3 EPSCs were recorded at -70mV and evoked at a 0.1Hz frequency of stimulation, whereas disynaptic IPSCs were recorded at +10mV (Fig. 4A and B). A minimal intensity of stimulation was used in order to elicit low amplitude EPSCs with a high level of failure, matching the well-described conditions of basal transmission at MF-CA3 PCs synapses (Lawrence et al., 2004; Marchal & Mulle, 2004) (Fig. S4B). At basal transmission (0.1Hz), the amplitude of evoked EPSCs and IPSCs (Fig. S4A and D) and the ratio between IPSCs and EPSCs (i.e. I/E ratio, Fig. 4C) were not different between genotypes. One typical feature of MF-CA3 PCs synapses is their ability to elicit robust low frequency facilitation (LFF) (Marneffe et al., 2025; Salin et al., 1996), by increasing the frequency of stimulation (e.g. from 0.1 to 1Hz). EPSCs and IPSCs evoked at a 1Hz stimulation frequency showed no difference in amplitudes between conditions, with equivalent I/E ratio (Fig. S4C and F and Fig. 4D, respectively). We then checked the levels of excitatory and inhibitory LFF for each genotype, and we observed a significant reduction in the 1Hz/0.1Hz ratio for both EPSCs and IPSCs in INT_ko compared to INT_ctl (Fig. 4E and F, mean ratio EPSCs: 4.2 ± 0.5 for INT_ctl (n=25), 2.6 ± 0.3 for INT_ko (n=20), unpaired T test: p = 0.0112; mean ratio IPSCs: 1.4 ± 0.2 for INT_ctl, 0.9 ± 0.1 for INT_ko, Mann-Whitney test: p = 0.0068). The pairwise comparison of IPSCs amplitude and charge from the recorded cells confirmed the absence of facilitation – or even synaptic depression in the cKO mice (Fig. 4G and Fig. S4H). Both the amplitude and the charge were significantly reduced at 1Hz compared to 0.1Hz stimulation frequency (mean amplitude IPSCs: 55.6 ± 13.6pA at 0.1Hz, 44.4 ± 10.7pA at 1Hz, Wilcoxon matched-pairs test: p = 0.0441; mean charge IPSCs: 72.7 ± 18.5pA/s at 0.1Hz, 56.4 ± 15.9pA/s at 1Hz, p = 0.0037, n=20). In control mice, there was no significant change of IPSCs amplitude and charge between rates of stimulation (Fig. 4H and Fig. S4I), showing globally no inhibitory low frequency facilitation in physiological condition at these CA3 INT-PC synapses. Additionally, the analysis of the distribution of IPSCs amplitude showed a significant shift towards smaller amplitude in APP/APLP cKO CA3 INTs at both 0.1Hz (basal transmission) and 1Hz frequency of stimulation (Fig. S4E and G, mean amplitude at 0.1Hz: 66.3 ± 2.6pA for INT_ctl (n=24), 51.8 ± 2.6pA for INT_ko (n=21), Kolmogorov-Smirnov test: p < 0.0001 with a bin width=10; mean amplitude at 1Hz: 77.5 ± 2.7pA for INT_ctl, 45.7 ± 1.9pA for INT_ko, Kolmogorov-Smirnov test: p < 0.0001, with a bin width=10).

**Figure 4.**
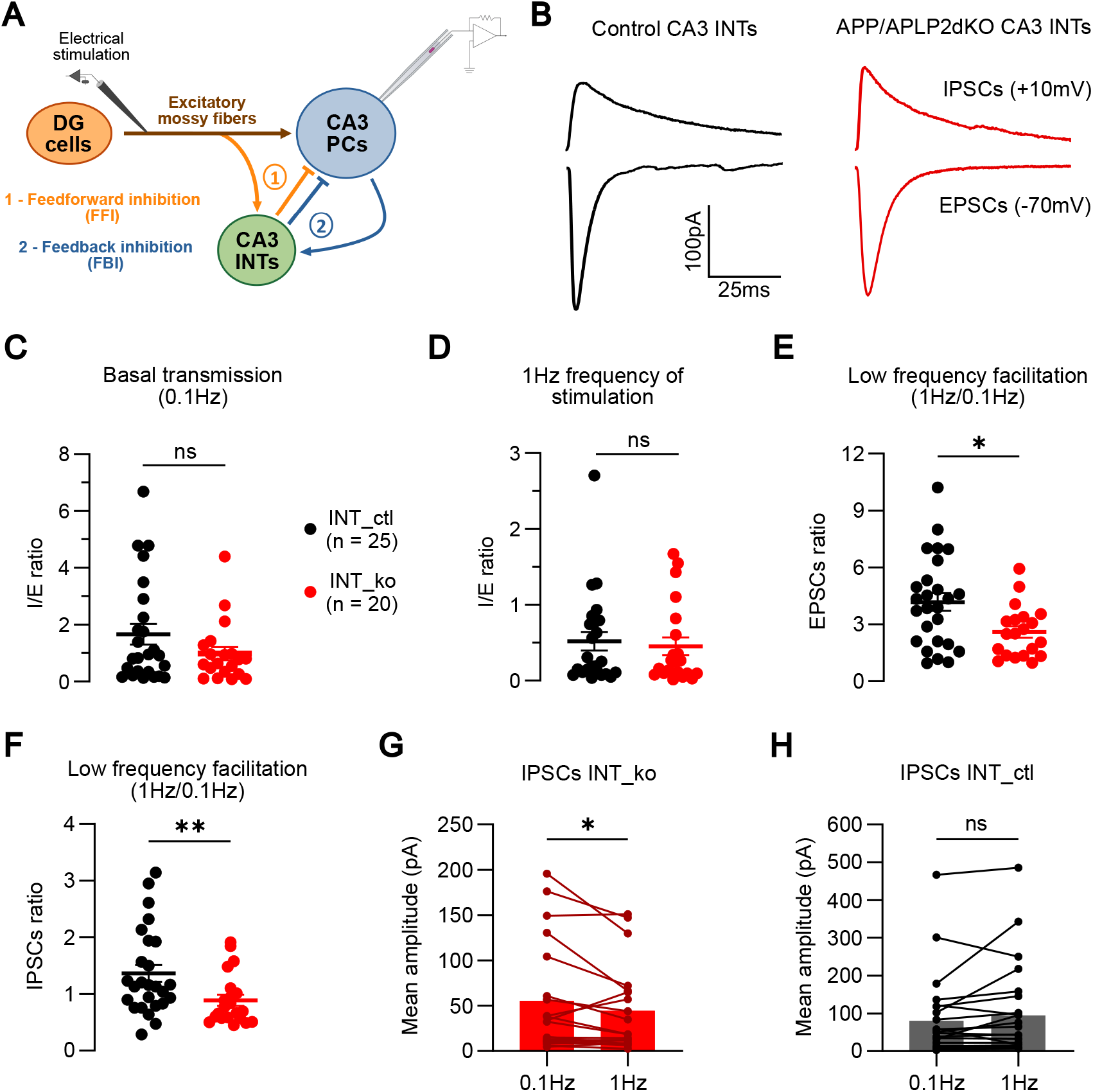
Basal transmission, low frequency facilitation and feedforward inhibition onto CA3 PC, in absence of APP/APLP2 in CA3 INTs. A. Schematic CA3 circuit with DG granule cells making excitatory connections through MFs onto both CA3 PCs and local INTs that send inhibitory projections onto PCs (i.e. feedforward inhibition). CA3 PCs are connected to CA3 INTs participating in feedback inhibition. Excitatory and inhibitory synaptic were recorded in CA3 PCs in the voltage-clamp mode. No drugs were added to the circulating ACSF for this set of experiments. B. Representative mean traces of evoked EPSCs and IPSCs recorded at negative (-70mV) and positive (+10mV) holding potential respectively, in INT_ctl (left - black) and in INT_ko (right - red). C and D. Mono- and di-synaptic responses were elicited by electrical stimulation of MFs. At 0.1Hz (basal transmission) and 1Hz frequency of stimulation, ratios between IPSCs and EPSCs (I/E ratio) are not altered in cKO condition compared to control (n=25 for INT_ctl; n=20 for INT_ko; Mann-Whitney test: not significant (ns)). However, 1Hz/0.1Hz ratios of both EPSCs (E) and IPSCs (F) are significantly reduced in INT_ko as compared to INT_ctl (mean ratio EPSCs: 4.2 ± 0.5 for INT_ctl, 2.6 ± 0.3 for INT_ko, unpaired T test: p = 0.0112; mean ratio IPSCs: 1.4 ± 0.2 for INT_ctl, 0.9 ± 0.1 for INT_ko, Mann-Whitney test: p = 0.0068). G. The pairwise comparison of recorded CA3 PCs in INT_ko showed a reduction of evoked IPSC amplitudes in INT_ko at 1Hz frequency of stimulation as compared to basal transmission (mean amplitude for INT_ko: 55.58 ± 13.65pA at 0.1Hz, 44.44 ± 10.73pA at 1Hz, Wilcoxon matched-pairs test: p = 0.0441). H. In control, there is no difference in the pairwise comparison of amplitude of IPSCs evoked by 0.1Hz and 1Hz frequency of stimulation.

Overall, these results could reflect an alteration of synaptic adaptability – allowing to adjust synaptic transmission in accordance to patterns of incoming activity – at inhibitory contacts onto CA3 PCs. Inhibitory synaptic depression is indeed observed when increasing the frequency of stimulation at MFs in the absence of APP/APLP2 in CA3 INTs. APP seems required for physiological mechanisms of synaptic release at higher frequency of stimulation in INTs. However, we do not have a clear explanation for the fact that excitatory presynaptic facilitation is also altered in the cKO group. The dual conditional deletion of APP and APLP2 impacts both LFF of excitatory transmission which is reduced but still present, as well as LFF of inhibitory transmission by inducing synaptic depression.

### *In vivo* study of the firing activity of CA3 circuits in CA3 INTs APP/APLP2 cKO mice

We next investigated whether the *ex vivo* functional perturbations depicted at the synaptic level impacted the activity of CA3 PCs and CA3 INTs at a circuit level. We performed *in vivo* silicon probe recordings of CA3 ensembles in head-fixed awake mice (APP^fl/fl^/APLP2^fl/fl^ mouse line or C57BL6/N for control), that received prior unilateral viral injection of the AAV9.mDLX-CRE-EGFP construct. After a 3-week period including implantation recovery and wheel apparatus habituation, head-fixed mice were confident on the wheel and were able to freely run with no signs of stress. We performed acute sessions of 30min duration during which both the brain activity (by extracellular electrophysiological recording using 64 channels – 4-shanks – silicone probe placed in CA3) and the locomotion on the wheel were recorded (Fig. 5A – left). After the recording session, mice were sacrificed for brain collection to later confirm viral expression and probe location in CA3 (Fig. 5A – right). From the raw electrophysiological signal, we extracted both neuronal spiking activity (obtained by spike sorting processing) and local field potential (LFP) oscillations (Fig. 5B). There is a strong relationship between the level of arousal and oscillatory activity in the brain (Buzsáki & Wang, 2012; Colgin, 2013). Using the LFP from CA3 stratum radiatum, we detected periods theta. While a large proportion of running co-occurred with theta, there was also theta detected in the absence of running. While there was large variability of the time spent in each state between animals (Fig. S5A, N=9 mice per genetic condition), there was no systematic difference between control and cKO mice. (Fig. S5B). The group comparison of the power spectrum (Fig. 5E) did not show significant differences in power in the following frequency ranges: theta (4-12Hz), gamma (20-50Hz), high gamma (50-100Hz), ripple (100-300Hz) (Fig. S5C).

**Figure 5.**
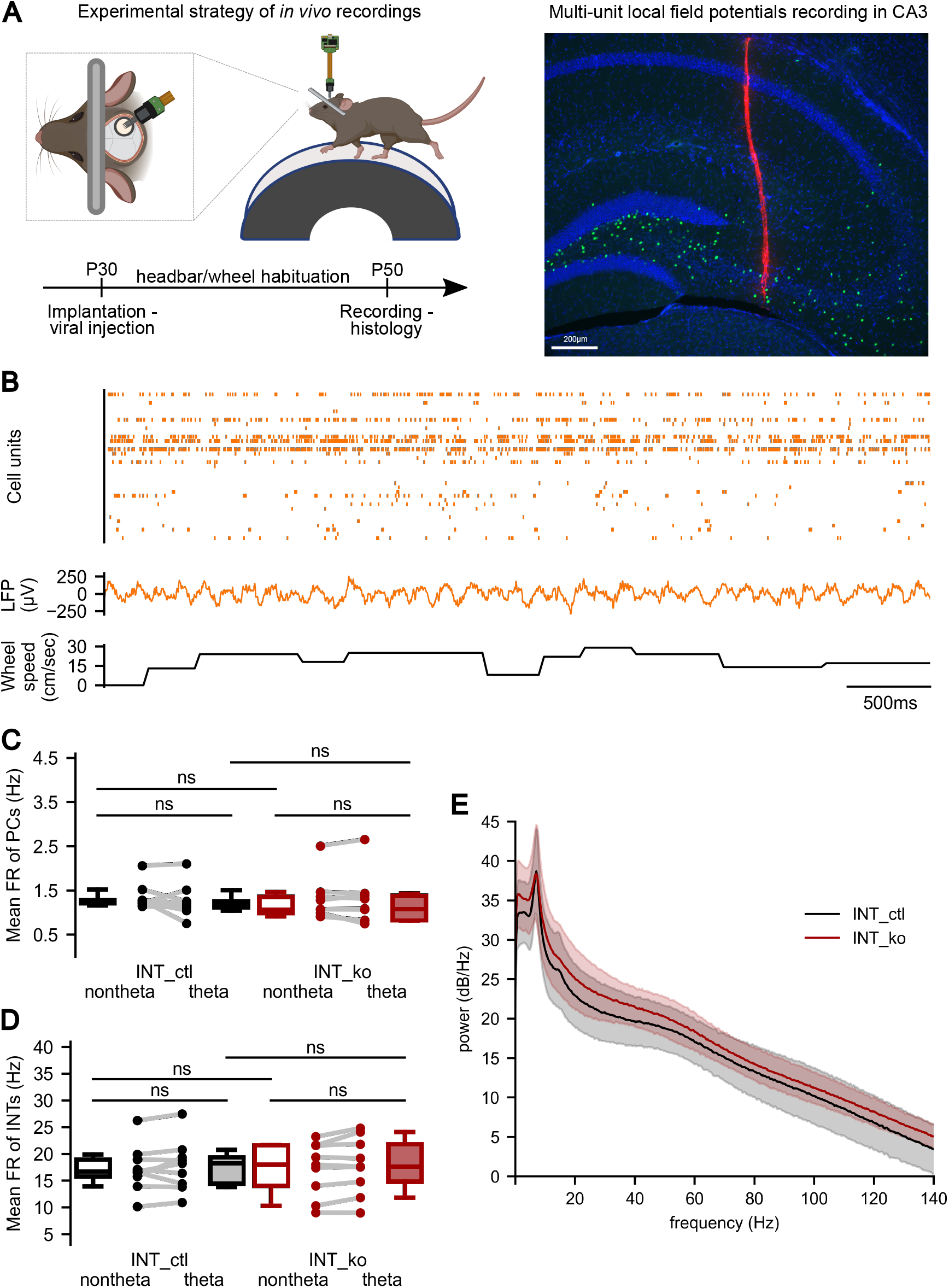
CA3 oscillations and neuronal dynamics in mice with selective deletion of APP/APLP2 in CA3 INTs. A. Left: experimental strategy for the in vivo silicone probe recordings in awake mice. At around 30 days post-natal (P30), unilateral CA3 injections of the AAV9.mDLX-Cre-EGFP viral construct and head bar implantation were performed in mice. 20 days later, acute in vivo extracellular electrophysiological recordings were made in CA3 in head-fixed awake mice allowed to run on a wheel. Both electrophysiological signals and animal mobility were recorded. Right: the brain collection allowed to check for probe location (DiI signal - red) and viral expression (EGFP - green) in CA3 cell layers (dapi - blue) on fixed slices. B. Raster plot highlighting the spiking activity of identified cell units (top), raw LFPs (middle) and locomotor activity(bottom) of one example recording in a control mouse. C and D. Boxplots of the mean firing rate (MFR) during non-theta and theta states for putative CA3 PCs and CA3 INTs, respectively (CA3 PCs MFR: 1.24 ± 0.27Hz (non-theta), 1.21 ± 0.35Hz (theta) for INT_ctl; 1.08 ± 0.47Hz (non-theta), 1.09 ± 0.55Hz (theta) for INT_ko; CA3 INTs MFR: 16.71 ± 4.16Hz (non-theta), 18.22 ± 4.50Hz (theta) for INT_ctl; 18.00 ± 4.77Hz (non-theta), 17.64 ± 5.05Hz (theta) for INT_ko. Values of the median ± standard deviation). There is no significant change of the MFR of CA3 PCs and INTs between non-theta and theta states in INT_ko. Wilcoxon signed-rank test: CA3 PCs for INT_ctl non-theta vs theta p = 0.25, INT_ko non-theta vs theta p = 0.57; CA3 INTs for INT_ctl non-theta vs theta p = 0.36, INT_ko non-theta vs theta p = 0.36. E. Average spectral density of CA3 oscillations from INT_ctl (grey) and INT_ko (red) recordings. Shaded regions represent the standard deviation.

The single units detected in the LFPs were classified into putative CA3 PCs and CA3 INTs based on waveform (i.e. trough-to-peak) and firing properties (i.e. mean firing rate, burst ratio and autocorrelogram width). The accuracy of segregation criteria of putative CA3 PCs and INTs was visually examined by comparing the mean waveform (Fig. S6A) and the cloud spreading on plots displaying parameters of detected PC and INT spiking units from the dataset (Fig. S6B). Average firing rates of putative CA3 PCs and INTs did not change between genetic conditions, neither during the theta or non-theta states (Fig. 5C-D).

In light of the strong recurrent connectivity between CA3 PCs thought to underpin ensemble formation, we quantified the level of the spike synchrony between pairs of putative PCs by taking the correlation coefficient of pairs of spike trains with 20ms binning. There was no difference in the cumulative distribution of the correlation coefficients of PC pairs between control and APP/APLP2 cKO mice (Fig. S7A, distribution of correlation coefficients: Mann Whitney U test, p = 1.00 INT_ctl *vs* INT_ko). A deeper analysis of CA3 PCs firing was made by looking at the number of population events (detected when the number of cells spiking together reached a defined threshold, see methods) in order to assess to what extent these events are influenced by genetic condition and brain state (Fig. S7B). The number of population events per second remained unchanged between non-theta and theta periods in both control and APP/APLP2 cKO mice (Fig. S7C). However, we found a significant reduction in the proportion of successive population events within 200ms windows, during theta periods as compared to non-theta in the control condition. This decrease was not present anymore in the APP/APLP2 cKO group (Fig. S7D, proportion of short inter-population-event-intervals (median ± MAD): 0.13 ± 0.12 (non-theta) 0.10 ± 0.10 (theta) for INT_ctl; 0.27 ± 0.12 (non-theta) 0.22 ± 0.15 (theta) for INT_ko; Wilcoxon signed-rank test: p = 0.031 for INT_ctl non-theta *vs* theta; p = 0.578 for INT_ko non-theta *vs* theta). Taken together, these results show that APP expression within CA3 INTs is not required to maintain a normal rate of cofiring within the CA3 PC network. Nevertheless, we found a defect in theta-dependent modulation of the temporal grouping of population events in APP/APLP2 cKO mice.

### Impaired theta-dependent modulation of CA3 PC spiking activity and theta-gamma cross-frequency coupling in CA3 INTs APP/APLP2 cKO mice

One main feature of the HPC is theta-phase spike modulation, wherein PCs and INTs firing is precisely controlled by the phase of theta, with the preferred phase depending on the cell identity. (Fernández-Ruiz et al., 2017). Moreoever, in CA1, it has been found that some populations of INTs, such as those expressing PV and SST, influence the theta-phase modulation of PCs (Royer et al., 2012). We explored whether theta spike modulation of excitatory PCs within CA3 were affected when local GABAergic INTs lack APP/APLP2, a manipulated population that should include perisomatic-targeting PV basket cells. The detected cycles of theta were time binned in order to calculate the circular distribution of spike counts by cell unit across theta cycle (from 0 to 360°) (Fig. 6A). In both INT_ctl and INT_ko, we identified “non theta phase-locked” and “theta phase-locked” CA3 PCs with fluctuating level of spike modulation (Fig. 6B). The average mean vector length (MVL – quantitative metric of the level of theta-spike modulation) showed a significant reduction in the INT_ko group (Fig. 6C, MVL (median ± MAD): 0.25 ± 0.05 for INT_ctl; 0.18 ± 0.06 for INT_ko, Mann-Whitney test, p = 0.042 INT_ctl *vs* INT_ko). However, the preferred phase of theta did not differ between genotypes (Fig. 6D-E, preferred phase given from 0 to 2π in radian). These results mean that CA3 PCs in APP/APLP2 cKO mice, with globally unchanged mean firing rates during theta period, fire with less temporal precision with respect to the theta cycle, whereas the theta phase during which PCs tend to preferentially discharge is not changed.

**Figure 6.**
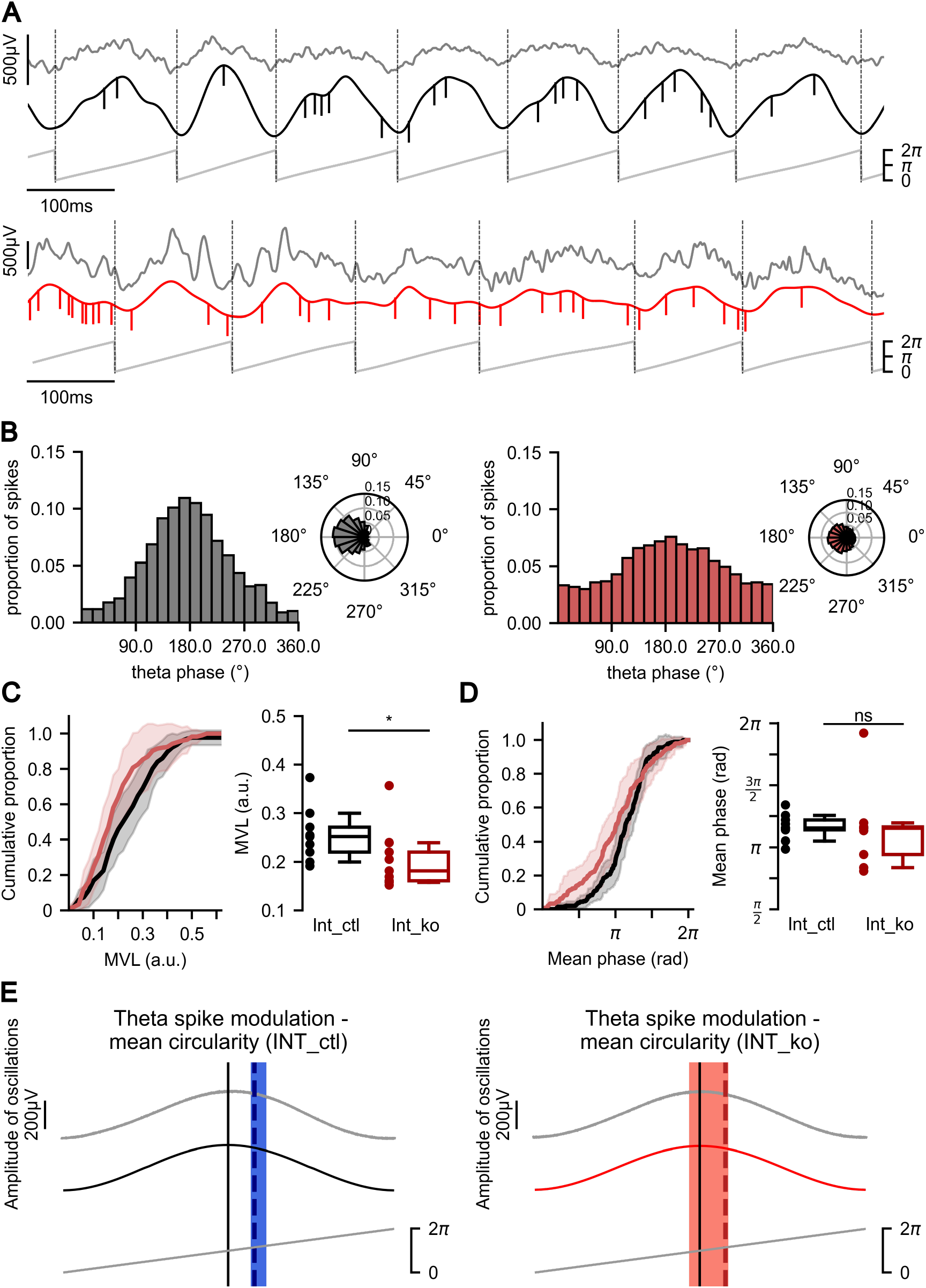
The deletion of APP/APLP2dKO in CA3 INTs reduces theta-spike modulation of CA3 PCs but does not modify theta mean phase. A. Example of recorded raw LFPs (dark grey), filtered theta oscillations, spiking activity (short vertical lines on top of theta show single spikes) and detected theta cycles (defined by dotted vertical lines - grey), in INT_ctl (top - dark) and INT_ko (bottom - red). B. Barplot of the distribution of spike proportions across theta, example cell unit in INT_ctl (left - darkgrey) and INT_ko (right - red). Circular plots (right side) illustrate the circular distribution of spikes across theta cycle, for each condition. C. Left: cumulative distributions of the mean vector length (MVL) of putative CA3 PCs in INT_ctl and INT_ko mice (back and red curves respectively). There is a significant reduction of theta-phase locking of PCs in INT_ko as compared to INT_ctl. Right: boxplot of the MVL with boxes defining the IQR and whiskers: 5th and 95th percentile. Individual points are MVL quantification by recording session. The MVL of each cell unit within a session has been inferred from the spike distribution across theta cycles (MVL: 0.25 ± 0.05 for INT_ctl; 0.18 ± 0.06 for INT_ko; Mann-Whitney U test, p = 0.042 INT_ctl vs INT_ko). Here and in the rest of the figure, data are expressed as median ± average absolute deviation from the median (MAD). D. Left: Cumulative distribution of the preferred phase of theta (i.e. mean phase in radians) showing no significant phase shift between conditions. Right: boxplot group comparison of theta mean phase (preferred phase, 3.63 ± 0.31 for INT_ctl; 3.62 ± 0.97 for INT_ko; Watson U-squared test, p = 0.773 INT_ctl vs INT_ko). E. Representative view of the mean prefered phase of CA3 PCs during theta cycle, in INT_ctl (left) and in INT_ko (right). In both conditions, excitatory cells preferentially discharge rapidly after theta peak, during the descending phase. The vertical dotted line represents the median and colored areas (blue - control and salmon - cKO) delimit Q1 (1st quartile) and Q3 (3rd quartile) of the mean circularity in each condition.

The theta-gamma phase-amplitude coupling is another remarkable oscillatory phenomenon of hippocampal circuits, where the amplitude of gamma oscillations is influenced by the phase of theta. Importantly, there is a strong implication of CA3 INTs in the generation of gamma oscillations within the HPC (Buzsáki & Wang, 2012; X. J. Wang & Buzsáki, 1996). The cross-frequency coupling increases the amplitude of gamma at the trough of theta phase in a consistent manner (Fig. 7A). We determined the level of gamma amplitude modulation during theta cycles by performing a cross-frequency analysis. The computed comodulograms across theta and gamma frequency ranges (from 1 to 16Hz and from 16 to 160Hz, respectively) were used to compare the averaged modulation index (MI) between genotypes (Fig. 7B). After averaging theta-gamma comodulograms in the 5-8Hz theta range, two dominant peaks of modulation at low and high gamma frequencies were identified (Fig. 7C). A significant decrease of the MI was observed for low gamma, but not high gamma, in APP/APLP2 cKO mice compared to control (Fig. 7D, MI in low-gamma frequencies (median ± MAD): 1.19e-3 ± 5.45e-4 for INT_ctl, 6.06e-4 ± 4.00e-4 for INT_ko, Mann-Whitney test: p = 0.042).. Overall, we observed no global change in the CA3 power spectrum (including theta and gamma frequency range) between control and experimental conditions. However, importantly, the precise study of theta spike modulation and theta-gamma cross-frequency coupling revealed that altered local inhibition impacts the oscillatory control of excitatory spiking dynamics and cross-frequency interactions.

**Figure 7.**
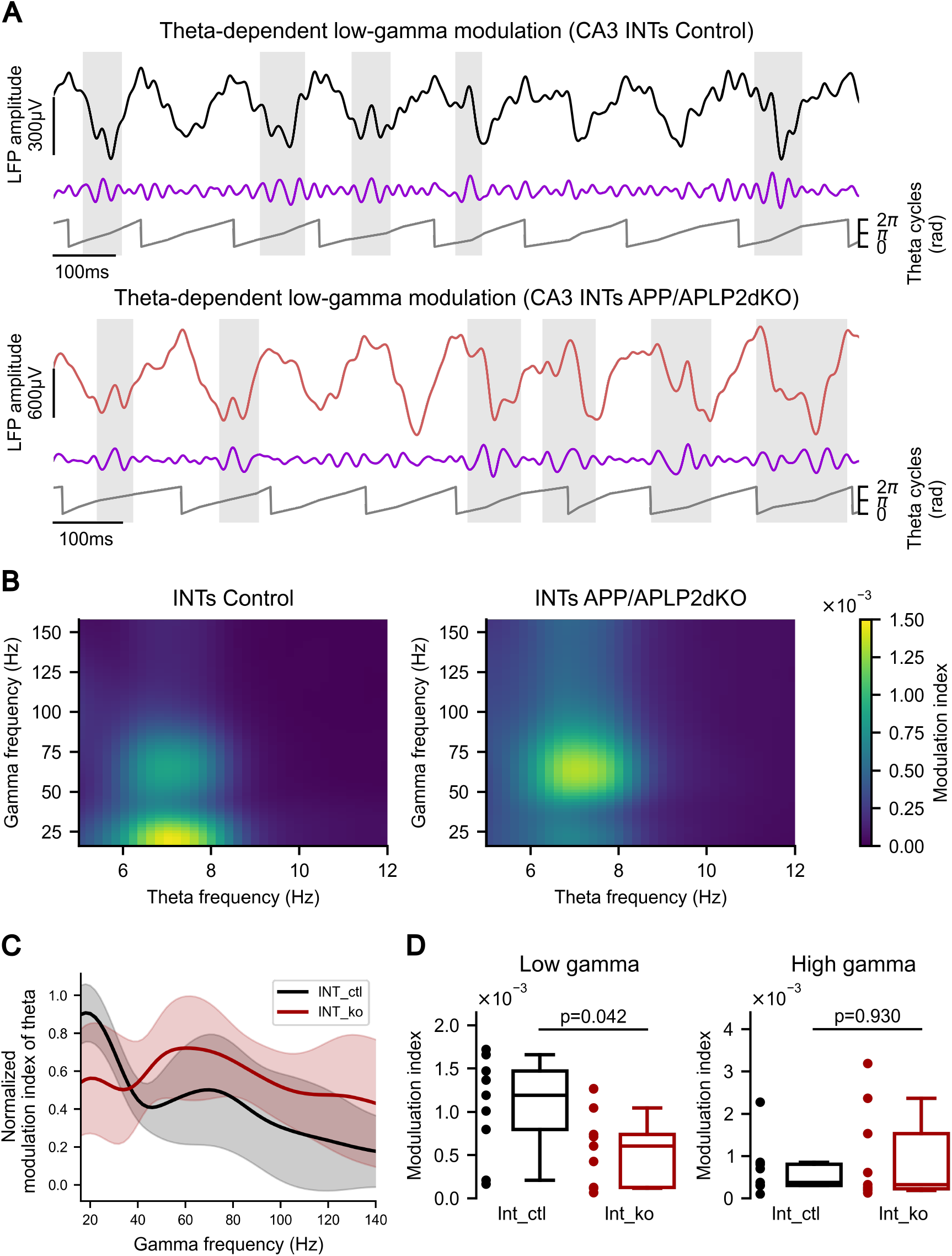
Specific perturbation of theta-dependent low-gamma amplitude modulation following APP/APLP2 deletion in CA3 INTs. A. Illustration of theta-gamma frequency-amplitude coupling in INT_ctl (top) and INT_ko mice (bottom). Example of raw LFPs (black/red traces), filtered low-gamma oscillations (blue) and theta cycles (dark grey). Light grey shaded areas highlight periods of low-gamma with increased amplitude, manually selected to illustrate gamma amplitude modulation by theta. B. Mean cross-correlograms in both conditions: INT_ctl (left) and INT_ko (right), highlighting two dominant hotspots with increased modulation index (MI) (i.e. low-gamma (16-40Hz) and high-gamma (40-110) respectively) within the range of theta (5-8Hz). C. Plot of the normalized modulation index of averaged theta within gamma frenquency range, highlighting two consistent peaks at low- and high-gamma frequencies in both INT_ctl and INT_ko conditions. Shaded areas represent the standard deviation in each group. D. Plots of the mean MI for theta across low-gamma (left) and high-gamma frequencies (right) showing a significant reduction of the MI for low-gamma in APP/APLP2 INT_ko mice. Boxes display the median, IQR (1st/3rd) and whiskers, the 5th and 95th percentiles. (MI in low-gamma range: 1.19e-03 ± 5.45e-04 for INT_ctl, 6.06e-04 ± 4.00e-04 for INT_ko; MI in high-gamma range: 3.80e-04 ± 6.18e-04 for INT_ctl, 3.27e-04 ± 1.06e-03 for INT_ko; Mann-Whitney U test: INT_ctl vs INT_ko, p = 0.042 for low-gamma and p = 0.930 for high-gamma). Data are expressed as median ± average absolute deviation from the median (MAD).

### Study of *in vivo* connectivity within the CA3 circuit in CA3 INTs APP/APLP2 cKO mice

The *ex vivo* results suggest a circuit reorganization at the synaptic level following APP/APLP2 deletion within CA3 INTs. We propose that under these conditions the cell connectivity within the CA3 circuit could be subsequently affected. In order to investigate this functional aspect *in vivo*, we performed a “pair-by-pair” analysis of the firing activity by extracting the cross-correlograms (CCGs) of the putative CA3 PCs and INTs. For pairs with a sufficient number of spikes, we were able to determine the type of connection between the “presynaptic” and the “postsynaptic” cell based on the firing of a "presynaptic" neuron with respect to the firing a putative connected "postsynaptic" neuron within a certain time window (1-5ms and 1.5-5ms for excitatory and inhibitory connections, respectively). The connection was considered as excitatory if the presynaptic cell significantly increased the number of APs in the downstream cell partner and considered as inhibitory for the opposite scenario – reduction of the downstream neuron firing (Fig. 8A, left and right CCG examples, respectively). The study of the number of connections (excitatory or inhibitory) per cell was performed on the entire dataset, including all the detected spiking units and ignoring the cell class (i.e. putative CA3 PCs, CA3 INTs and unknown cells). No differences were found in the number of downstream excitatory partners per cell and in the number of inhibitory partners per cell between INT_ctl and INT_ko genotypes (Fig. 8B and C). We then analyzed the area under the curve of the CCGs (used as a proxy to evaluate the strength of connection) within the critical time window of firing. The global normalized strength of excitatory connections and of inhibitory connections was not modified between INT_ctl and INT_ko (Fig. 8D and E, respectively). In order to increase our sample size for consistent statistics and at the condition that the CCG shape was describing an inhibitory connection, the cell units previously detected as “unknown” (excluded from putative CA3 PC and not entirely fitting the criteria for CA3 INT during cell classification processing) were categorized as INT cell type and used for further connection strength analysis. Interestingly, the investigation of the strength of connection while taking into account the cell class (INT or PYR) of the presynaptic and the postsynaptic neuron and the state (non-theta *versus* theta) revealed significant differences relative to state and conditions. First, we observed a significant increase of the synaptic strength during theta for INT onto INT type of connections (inhibitory synapses), compared to non-theta (Fig. 8F, INT-INT strength of connection: 0.51 ± 0.14 INT_ctl (non-theta), 1.26 ± 0.27 INT_ctl (theta), 0.43 ± 0.13 INT_ko (non-theta), 0.68 ± 0.24 INT_ko (theta); Mann Whitney U test: p = 0.39 INT_ctl (non-theta) *vs* INT_ko (non-theta), p = 0.015 INT_ctl (theta) *vs* INT_ko (theta); Wilcoxon signed-rank test: p = 0.0312 for INT_ctl non-theta *vs* theta; p = 0.0312 for INT_ko non-theta *vs* theta). A p_value_ correction for false discovery rate (FDR) was applied by using Benjamini/Hochberg and Benjamini/Yekutieli methods on independent values (state comparison) and negatively correlated values (genotype comparison), respectively (corrected p_value_ INT_ctl (non-theta) *vs* INT_ctl (theta): p = 0.047; corrected p_value_ INT_ko (non-theta) *vs* INT_ko (theta): p = 0.047; corrected p_value_ non-theta INT_ctl *vs* INT_ko: p = 0.39; corrected p_value_ theta INT_ctl *vs* INT_ko: p = 0.030). Secondly, the connection strength during theta periods was significantly reduced in INT_ko compared to INT_ctl, specifically for that synapse type (INT-INT connection). The analysis of other types of connections (i.e. INT-PYR and PYR-INT) showed no effects of APP/APLP2 removal within CA3 INTs on the strength of connections, and of the state (Fig. 8G and H). However, the analysis highlighted a consistent but not significant augmentation of the strength of connection in both genotypes during theta compared to non-theta, primarily for PYR-INT connections (Fig. 8H). Finally, the detailed analysis of the firing behavior of interconnected cell units highlighted differences in our dataset attributed to specific types of detected neuronal connections, where inhibitory synapses initiated by INTs contacting other INTs are the most susceptible to be affected by the conditional suppression of APP and APLP2 in INTs. In fact, during theta activity when neuronal ensembles are very likely to synchronize and discharge together, the synaptic properties of local INTs to elicit indirect disinhibition of CA3 PCs via the inhibition of nearby INTs appear to be weakened. Such functional modifications potentially relate to the previous observations made at the synaptic level, suggesting a structural reorganization of inhibitory inputs in CA3 circuits.

**Figure 8.**
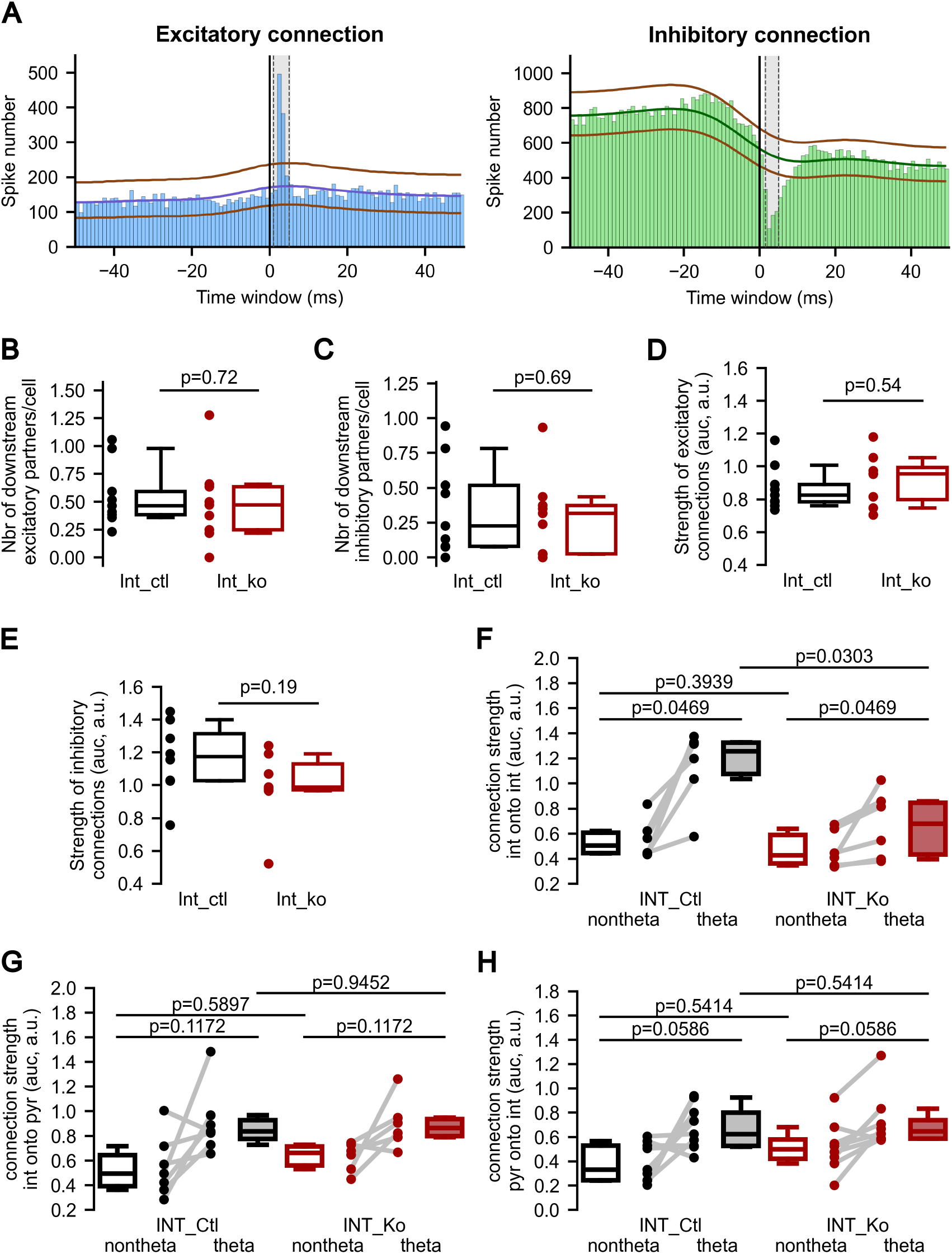
In vivo cell connectivity of populations of principal cells and interneurons within CA3 following APP/APLP2 deletion in CA3 INTs. A. Left: illustration of pairs of connected cells forming either an excitatory connection (cross-correlogram example where postsynaptic cell spiking activity is significantly increased under the control of presynaptic firing). Right: inhibitory connection formed by a pair of cells where postsynaptic cell spiking activity is significantly decreased). Excitatory and inhibitory connections are made by an upstream excitatory and inhibitory presynaptic partner, respectively. The black vertical line highlights T0 (spike time of the presynaptic cell) and grey dotted lines define the time window of interest (1.5-5ms) to define a connection. Horizontal brown lines mark upper and lower limits to reach connectivity threshold, and blue and green lines are the fitted curves of the gaussian filter applied to the bin values for excitatory and inhibitory connections, respectively. B. Mean number of downstream excitatory partners per cell for INT_ctl and INT_ko. C. Mean number of downstream inhibitory partners per cell. Values obtained for each session by averaging the number of detected outgoing connections between pairs cells, either excitatory or inhibitory. Boxes indicate the 1st, 2nd and 3rd quartiles and the whiskers are set to the 5th and 95th percentiles of the values. Plots include individual data points in each condition (statistics: Mann-Whitney U test: p > 0.05 (not significant)). D. Normalized strength of excitatory connections for INT_ctl and INT_ko mice. E. Normalized strength of inhibitory connections. The area under the curve was used as a proxy to evaluate the strength of connection. Values of connection strength were normalized to the bound (upper for excitatory connections or lower for inhibitory ones). F. A deeper analysis of the strength of connections highlight a significant reduction during theta state for INT_ko as compared to INT_ctl mice, specifically for INT-INT type of inhibitory connections, as well as a consistent increase of the strength of connection in both INT_ctl and INT_ko during theta compared to non-theta (connection strength: 0.51 ± 0.14 INT_ctl (non-theta), 1.26 ± 0.27 INT_ctl (theta), 0.43 ± 0.13 INT_ko (non-theta), 0.68 ± 0.24 INT_ko (theta); corrected p_value_ INT_ctl (non-theta) vs INT_ctl (theta): p = 0.047; corrected p_value_ INT_ko (non-theta) vs INT_ko (theta): p = 0.047; corrected p_value_ INT_ctl (non-theta) vs INT_ko (non-theta): p = 0.39; corrected p_value_ INT_ctl (theta) vs INT_ko (theta): p = 0.030). G and H. No significant modification of the synaptic strength is observed for both state and group conditions for other types of connections.

## Discussion

Even though APP is ubiquitously expressed in neurons, its cell-specific physiological functions remain incompletely understood. In this study, we have explored the physiological role of APP in GABAergic INTs which have received relatively less attention, as compared to excitatory neurons and synapses (Dunot et al., 2023; Müller et al., 2017). We have targeted the hippocampal CA3 region, which plays a major role in episodic memory encoding and retrieval, cognitive processes which are severely affected early during the course of AD. GABAergic INTs in CA3 were targeted using a viral construct expressing Cre under the control of mDLX, to delete both APP and APLP2. Single APP KO results in a subtle phenotype and lack major synaptic alterations found in APP/APLP2dKO mice (Fanutza et al., 2015). Inversely, constitutive APP/APLP2dKO mice rapidly die after birth due to morphological synaptic deficiency (Korte et al., 2012; P. Wang et al., 2005). The full extent of APP function is thought to be compensated by other members of the APP family, in particular by APLP2 (Hick et al., 2015), justifying the use of a genetic model with deletions of both APP and APLP2. By performing *ex vivo* and *in vivo* electrophysiology experiments, we show that the deletion of APP/APL2 in CA3 GABAergic INTs only has a minor impact on the morphological properties and intrinsic excitability of GABAergic INTs, however it has a significant impact on the inhibitory coverage of CA3 PCs. *In vivo*, this translates into a dysregulation of functional connectivity between GABAergic INTs which may contribute to the changes observed in the oscillatory control of spiking dynamics and inter-frequency rhythmic interactions.

A previous study has taken advantage of conditional APP/APLP2 double knockout mice in all GABAergic forebrain INTs by crossing APP^fl^/APLP2^fl^ mice with DlxCre mice (Mehr et al., 2020). In hippocampal slices of DlxCre cDKO mice, deficits in excitation/inhibition balance, with a relative excess in synaptic inhibition, and reduced firing rate of CA1 PCs were observed, albeit with no change in spontaneous IPSCs (Mehr et al., 2020). This was accompanied with deficits in hippocampus-dependent memory tasks (Mehr et al., 2020). Interestingly, the morphological complexity of CA1 INTs was increased, selectively for calbindin-positive neurons (Mehr et al., 2020). Overall, this study concluded that APP/APLP2 in GABAergic INTs play distinctive roles in hippocampal function, which may affect the activity of local circuits. It should be considered however that the dysregulation of neuronal excitability, and its behavioral consequences, may arise, at least in part, from a developmental effect of APP on neuronal morphology and synaptic connectivity (Müller et al., 2017). More recently, it was shown that the conditional deletion of APP, APLP1 and APLP2 in all GABAergic neurons resulted in increased GABAergic synaptic responses in hippocampal CA1 PCs without any change in short-term plasticity (Lee et al., 2026). Our study takes a step further by exploring the link between the properties of CA3 INTs *ex vivo* and local CA3 circuit activity *in vivo*, in conditions of APP/APLP2 deletion after development has taken place.

A question which has remained unanswered in these previous studies was the impact of APP/APLP2 deletion on the intrinsic excitability of local GABAergic INTs. This question is important in view of the rising evidence that hyperexcitability linked to excitation/inhibition imbalance is an early pathophysiological hallmark of AD (Frere & Slutsky, 2018; Palop & Mucke, 2016; Xu et al., 2020; Zott et al., 2018), which has been observed both in animal models and AD patients with Mild Cognitive Impaitment (MCI) (Bakker et al., 2012). GABAergic INTs are thus central players in the pathophysiology of AD, and as such it is important to explore the physiological role of APP in this cell population. We have found only minor changes in the morphological and functional intrinsic properties when considering all GABAergic INTs, fluorescently labeled using the AAV9.mDLX-CRE-EGFP viral construct. In order to analyze whether a selective subpopulation of INTs was impacted by the deletion of APP/APLP2, we performed a dimensionality reduction analysis based on the electrophysiological parameters of recorded INTs. Three clusters of cells with markedly distinct functional properties represented the high heterogeneity of INTs in CA3 (Pelkey et al., 2017; Szabadics & Soltesz, 2009; Tzilivaki et al., 2023). For one of the clusters, an increase in input resistance was observed for the INT_ko group, whereas the deletion of APP/APLP2 had no effect on any other electrophysiological or morphological parameters. Overall, our results indicate that APP and APLP2 do not play a major role in shaping the intrinsic excitability of GABAergic INTs. This contrasts with reports of impaired intrinsic excitability in GABAergic INTs in different mouse models of APP-related pathology (Goettemoeller et al., 2024; Hijazi et al., 2020, 2023; Olah et al., 2022; Petrache et al., 2019; Verret et al., 2012), suggesting that these alterations may be due to the overexpression of hAPP, rather than to variations in the physiological function of APP.

Numerous studies have implicated APP (and APLP2) in synaptic transmission, in both pre- and postsynaptic mechanisms, and mostly at excitatory synapses (Barthet & Mulle, 2020; Dunot et al., 2023; Müller et al., 2017). However, the constitutive removal of APP/APLP2 or APP/APLP1/APLP2 in GABAergic INTs affects neither the frequency nor the amplitude of mIPSCs in CA1 PCs (Lee et al., 2026; Mehr et al., 2020). In contrast, we observe a marked change in the distribution of spontaneous and miniature IPSCs when recording CA3 PCs in APP/APLP2 cKO mice, with a larger proportion of IPSCs with higher amplitude and faster rise time. The localization of GABAergic synaptic contacts onto principal neurons is highly heterogeneous (soma, axon initial segment, dendritic shafts and spines), supporting different modes of control of the postsynaptic neuron (Kubota et al., 2016; Miles et al., 1996), with a specific temporal activation profile critically contributing to brain oscillations (Cupolillo et al., 2024; Tzilivaki et al., 2023). Our results are consistent with a relative increase in the number and/or efficacy of perisomatic synapses, characterized by a fast onset and high amplitude of IPSCs (Booker & Vida, 2018; Kubota et al., 2016). Basket cells which represent the major population of perisomatic-targeting INTs comprise PV-positive and CCK-positive INTs (Booker & Vida, 2018; Kubota et al., 2016). PV-positive basket cells, which provide rapid and effective synaptic inhibition as compared to CCK-positive basket cells (Kohus et al., 2016), thus appear as a potential candidate implicated in the phenotype observed in the absence of APP/APLP2. Additional experiments that include paired recordings of PV-positive INTs and CA3 PCs should help in understanding the mechanisms by which the properties of perisomatic GABAergic synapses change in APP/APLP2 cKO condition. APP/APLP2 removal may preferentially affect neuronal activity of PV-positive INTs rather than of other subpopulations of INTs. The observed modifications may also rely on presynaptic mechanisms by increasing the probability of GABA release at synapses between PV-positive INTs and CA3 PCs, which is already high in control conditions (Kohus et al., 2016). Conversely, an increase of the morphological coverage of CA3 PCs by axons of PV-positive INTs, in the absence of functional changes in synaptic mechanisms would also be consistent with the phenotype described.

Our results point to a functional reorganization of inhibition of CA3 PCs which may involve PV-positive INTs following APP/APLP2 deletion. Multiple studies have reported that PV-positive INTs are central to the dysfunction of neural circuits and related cognitive decline in AD mouse models (Goettemoeller et al., 2024; Hijazi et al., 2023, 2023; Olah et al., 2022; Petrache et al., 2019). The impairment of PV neurons in hAPP mice is accompanied with a decreased expression of the Nav1.1 sodium channel subunit and a similar decrease is reported in post-mortem brain tissue of AD patients, thought to cause alterations of oscillatory activity (Verret et al., 2012). Consistently, the impaired function of PV-positive INTs in mouse models of AD results in deficient inhibitory control, changes in oscillatory activity, and overall increased excitation (Hijazi et al., 2023). Confirming previous studies (B. Wang et al., 2014), we have observed that APP is expressed in all GABAergic INTs in CA3, and not selectively in PV-positive INTs. Whether APP plays a role in the potential vulnerability of PV positive INTs in models of AD is currently unclear, albeit our results do not favor a distinctive role of APP/APLP2 in intrinsic excitability properties in any INT subpopulation, including PV positive INTs.

In order to search for a correlation between the reorganization of inhibition in CA3 found *ex vivo*, and neuronal activity and connectivity in relation to oscillations, we performed *in vivo* electrophysiological recordings of CA3 region in awake mice with deletion of APP/APL2 in GABAergic INTs. Many studies have provided evidence that GABAergic INTs are crucial for the generation of brain oscillations in the HPC and for network synchronization, two neuronal mechanisms required for cognitive functions such as learning and memory (Bartos et al., 2002, 2007; Bocchio et al., 2024; Eichenbaum, 2017; Vida et al., 2006). Alterations in circuit dynamics related to impairment of inhibition have been clearly demonstrated in mouse models of AD and in patients (Carello-Collar et al., 2023; Ebrahimi et al., 2025; Govindpani et al., 2017; Lanctot et al., 2007; Palop et al., 2007; Palop & Mucke, 2016; Verret et al., 2012; Viana da Silva et al., 2023; Wu et al., 2014). However, the underlying inhibitory mechanisms need to be further characterized *in vivo*. Also, it remains unclear whether selected subpopulations of INTs contribute more significantly to circuit dysfunction associated with AD pathophysiology or whether it is caused by a global defect of inhibition which does not rely on a specific cell-type. The two main subtypes of basket cells, PV-positive and CCK-positive perisomatic targeting INTs, have different contribution to network oscillations (Rozov et al., 2024). Within CA3, PV-positive basket cells, making inhibitory contacts onto the soma and proximal dendrites of principal neurons, preserve incoming activity patterns and contribute to oscillatory activity and network synchrony by displaying fast, reliable and temporally sharp synaptic control of excitatory neurons (Freund & Katona, 2007; Tukker et al., 2013). In contrast, CCK-positive basket cells, which are highly diversified, are globally less coupled to fast rhythmic activity and provide more imprecisely timed inhibitory outputs rather supporting neuronal adaptation and synaptic plasticity through disinhibition processes (Bartos & Elgueta, 2012; Galarreta & Hestrin, 2001; Hu et al., 2010; Lasztóczi et al., 2011; Park et al., 2020). In addition, as largely described in CA1, INTs display distinct temporal patterns of firing during theta oscillations, being either independent or tightly controlled by theta rhythms (Czurkó et al., 2011; Klausberger & Somogyi, 2008; Lasztóczi et al., 2011; Royer et al., 2012; Viney et al., 2013).

Silicon probe electrophysiological recordings in mice lacking APP/APLP2 in CA3 INTs did not reveal any major changes in CA3 power spectrum (including in the theta and gamma frequency ranges) or in the overall spiking activity of CA3 principal neurons between control and experimental conditions. However, we observed a reduction of theta-low gamma cross-frequency coupling in APP/APLP2 cKO mice. Furthermore, we highlighted a reduction of theta spike modulation of CA3 PCs in these mice. Theta spike-phase modulation is an efficient activity mechanism that promotes cell-specific discharge with temporal rhythmicity. Interestingly, in the HPC, theta is strongly associated with mobility and “on-line” brain state during which specific inhibitory circuits are activated (Wester & McBain, 2014). Indeed, there is a positive but not exclusive correlation between locomotion and neuronal activation of SST-expressing and PV-expressing CA1 INTs (Arriaga & Han, 2017). We found equivalent proportions of excitatory population events in both genetic conditions in CA3, regardless of the brain state. However, we observed that the modulation of the number of “bursty” population events of CA3 PCs, occurring during theta, is no longer present in mice lacking APP/APLP2 in CA3 INTs. It could reflect a perturbation of the inhibitory control exerted on excitatory ensembles during theta, possibly relying on SST-positive INTs (Royer et al., 2012). We also observed *in vivo* a reduction of the connection strength during theta in INT_ko animals, specifically for pairs of INTs, although the inhibitory synaptic strength appeared globally unchanged between groups.

Neuronal populations of different cell types do not equally contribute to hippocampal rhythms. Moreover, oscillatory-dependent CA3 PC firing most likely relies on a robust functional interconnection between INTs. Based on our findings in slices, PV-positive INTs are most susceptible to driving reorganization of inhibitory inputs onto CA3 PCs in APP/APLP2 cKO mice. The observed change in inhibitory coverage in CA3 PCs would likely enhance the modulation of CA3 PC spiking during theta, via powerful and precise temporal control driven by PV-positive INTs (Amilhon et al., 2015; Freund & Katona, 2007; Royer et al., 2012; Varga et al., 2012). However, *in vivo* recordings rather reveal a decreased theta-dependent modulation of excitatory spike timing in APP/APLP2 cKO mice. It would be interesting to explore whether the connection strength between INTs, modulated by theta, could play a role in the reduction of theta-spike modulation of CA3 PCs.

INTs are critical elements in regulating information processing by adapting and undergoing plastic modifications to maintain a coherent control of PCs firing (Topolnik & Tamboli, 2022). There are for instance subtype-specific and activity dependent changes of INT dynamics that occur during precise oscillatory patterns, as recently shown during sharp-wave ripples (SWRs) in CA3/CA2 circuits (Vancura et al., 2023). Given the tight functional relationship between GABAergic inhibition, and theta and gamma activity, we propose that in the absence of APP/APLP2 in CA3 INTs, there is a specific loss of plastic properties of INT-INT and INT-PC synapses, consequently perturbing the control of oscillatory dynamics and of firing of PC ensembles in CA3. Interestingly, in the context of AD where there is a disruption of APP functions, similar perturbations of the oscillation component have been highlighted. In hAPP AD mouse model, modifications of PV INT neuronal properties lead to a reduction of gamma oscillations and network synchrony (Verret et al., 2012). Alongside, localized expression of hAPP in CA3 excitatory inputs perturbs cellular to circuit interface, theta oscillation amplitude and precise temporal dynamics of downstream CA1 PC spiking relative to theta (Viana da Silva et al., 2023). Lastly, *in vivo* optogenetic or chemogenetic manipulation of PV-positive INTs (possibly contributing to the APP/APLP2 cKO phenotype) in AD transgenic mice, can prevent or rescue impaired brain oscillations, like slow-gamma, and memory deficits (Etter et al., 2019; Hijazi et al., 2020).

In summary, our combined *ex vivo* and *in vivo* experiments show that the selective deletion of APP (and APLP2) in INTs does not affect directly their intrinsic excitability but leads to a reorganization of inhibitory inputs onto CA3 PCs, affects connectivity strength between INTs in a theta-dependent manner, and finally impairs theta-gamma coupling. In future experiments, it would be interesting to understand whether these mechanisms are functionally related or occur independently, and whether they are caused by deficits in specific INT populations. Overall, APP and APLP2 control inhibitory and excitatory synaptic functions through complementary mechanisms in principal neurons and interneurons. Our study suggests an important physiological role of APP and APLP2 expressed in INTs, in controlling local circuit activity in reference to theta-gamma oscillations, brain mechanisms that are compromised in AD.

## Acknowledgements

This work was supported by the Centre National de la Recherche Scientifique (CNRS), the University of Bordeaux, the French ANR (GenSynAlz and AD-PLCG2 grants to C.M), the Fondation Vaincre Alzheimer (PhD fellowship to AE). We thank Gaël Barthet for his invaluable initial input on the project and for Alexandre Favereaux for the thoughtful discussions on the clustering analysis. We thank Cindie Leteneur for her help for the first patch-clamp experiments. We thank the Bordeaux Imaging Center (BIC), a service unit of the CNRS-Inserm and Bordeaux University, member of the national infrastructure France BioImaging, the Pôle in Vivo for their continuous help with hosting the mice, and the AAV vector facility for providing AAV particles.

## Author contributions

A.E, A.MdS and C.M conceived the study and designed the experiments. A.E and A.MdS performed the *in vitro* electrophysiological experiments. A.E performed the *in vivo* recordings. A.E and A.K processed and analyzed the *in vivo* data. E.G and S.D helped in the design and tests of the viral constructs. All authors and acknowledged collaborators took part in the design of experimental protocols and data interpretation. A.E and C.M wrote the manuscript.

## Bibliography

Amilhon, B., Huh, C. Y. L., Manseau, F., Ducharme, G., Nichol, H., Adamantidis, A., & Williams, S. (2015). Parvalbumin Interneurons of Hippocampus Tune Population Activity at Theta Frequency. Neuron, 86(5), 1277– 1289. 10.1016/j.neuron.2015.05.027

Arriaga, M., & Han, E. B. (2017). Dedicated Hippocampal Inhibitory Networks for Locomotion and Immobility. The Journal of Neuroscience: The Official Journal of the Society for Neuroscience, 37(38), 9222–9238. 10.1523/JNEUROSCI.1076-17.2017

Atallah, B. V., & Scanziani, M. (2009). Instantaneous Modulation of Gamma Oscillation Frequency by Balancing Excitation with Inhibition. Neuron, 62(4), 566–577. 10.1016/j.neuron.2009.04.027

Bakker, A., Krauss, G. L., Albert, M. S., Speck, C. L., Jones, L. R., Stark, C. E., Yassa, M. A., Bassett, S. S., Shelton, A. L., & Gallagher, M. (2012). Reduction of hippocampal hyperactivity improves cognition in amnestic mild cognitive impairment. Neuron, 74(3), 467–474. 10.1016/j.neuron.2012.03.023

Barthet, G., & Mulle, C. (2020). Presynaptic failure in Alzheimer’s disease. Progress in Neurobiology, 194, 101801. 10.1016/j.pneurobio.2020.101801

Bartos, M., & Elgueta, C. (2012). Functional characteristics of parvalbumin- and cholecystokinin-expressing basket cells. The Journal of Physiology, 590(4), 669–681. 10.1113/jphysiol.2011.226175

Bartos, M., Vida, I., Frotscher, M., Meyer, A., Monyer, H., Geiger, J. R. P., & Jonas, P. (2002). Fast synaptic inhibition promotes synchronized gamma oscillations in hippocampal interneuron networks. Proceedings of the National Academy of Sciences, 99(20), 13222–13227. 10.1073/pnas.192233099

Bartos, M., Vida, I., & Jonas, P. (2007). Synaptic mechanisms of synchronized gamma oscillations in inhibitory interneuron networks. Nature Reviews Neuroscience, 8(1), 45–56. 10.1038/nrn2044

Bateman, R. J., Aisen, P. S., De Strooper, B., Fox, N. C., Lemere, C. A., Ringman, J. M., Salloway, S., Sperling, R. A., Windisch, M., & Xiong, C. (2011). Autosomal-dominant Alzheimer’s disease: A review and proposal for the prevention of Alzheimer’s disease. Alzheimer’s Research & Therapy, 3(1), 1. 10.1186/alzrt59

Bocchio, M., Vorobyev, A., Sadeh, S., Brustlein, S., Dard, R., Reichinnek, S., Emiliani, V., Baude, A., Clopath, C., & Cossart, R. (2024). Functional networks of inhibitory neurons orchestrate synchrony in the hippocampus. PLoS Biology, 22(10), e3002837. 10.1371/journal.pbio.3002837

Booker, S. A., & Vida, I. (2018). Morphological diversity and connectivity of hippocampal interneurons. Cell and Tissue Research, 373(3), 619–641. 10.1007/s00441-018-2882-2

Buhl, E. H., Halasy, K., & Somogyi, P. (1994). Diverse sources of hippocampal unitary inhibitory postsynaptic potentials and the number of synaptic release sites. Nature, 368(6474), 823–828. 10.1038/368823a0

Burns, A. P., Duran, M. I., Fortel, I., Lazarov, O., Zhan, L., Bendlin, B., & Leow, A. (2026). Excitation–inhibition homeostasis in Alzheimer’s disease: A selective multiscale review of mechanisms, sex differences, and therapeutic opportunities. Neuropsychopharmacology, 1–17. 10.1038/s41386-026-02334-0

Busche, M. A., Kekuš, M., Adelsberger, H., Noda, T., Förstl, H., Nelken, I., & Konnerth, A. (2015). Rescue of long-range circuit dysfunction in Alzheimer’s disease models. Nature Neuroscience, 18(11), 1623–1630. 10.1038/nn.4137

Busche, M. A., & Konnerth, A. (2016). Impairments of neural circuit function in Alzheimer’s disease. Philosophical Transactions of the Royal Society B: Biological Sciences, 371(1700), 20150429. 10.1098/rstb.2015.0429

Buzsáki, G., & Wang, X.-J. (2012). Mechanisms of gamma oscillations. Annual Review of Neuroscience, 35, 203– 225. 10.1146/annurev-neuro-062111-150444

Carello-Collar, G., Bellaver, B., Ferreira, P. C. L., Ferrari-Souza, J. P., Ramos, V. G., Therriault, J., Tissot, C., De Bastiani, M. A., Soares, C., Pascoal, T. A., Rosa-Neto, P., Souza, D. O., & Zimmer, E. R. (2023). The GABAergic system in Alzheimer’s disease: A systematic review with meta-analysis. Molecular Psychiatry, 28(12), 5025–5036. 10.1038/s41380-023-02140-w

Chamberland, S., Grant, G., Machold, R., Nebet, E. R., Tian, G., Stich, J., Hanani, M., Kullander, K., & Tsien, R. W. (2024). Functional specialization of hippocampal somatostatin-expressing interneurons. Proceedings of the National Academy of Sciences of the United States of America, 121(17), e2306382121. 10.1073/pnas.2306382121

Colgin, L. L. (2013). Mechanisms and Functions of Theta Rhythms. Annual Review of Neuroscience, 36, 295– 312. 10.1146/annurev-neuro-062012-170330

Csicsvari, J., Hirase, H., Czurkó, A., Mamiya, A., & Buzsáki, G. (1999). Oscillatory Coupling of Hippocampal Pyramidal Cells and Interneurons in the Behaving Rat. Journal of Neuroscience, 19(1), 274–287. 10.1523/JNEUROSCI.19-01-00274.1999

Cupolillo, D., Regio, V., & Barberis, A. (2024). Synaptic microarchitecture: The role of spatial interplay between excitatory and inhibitory inputs in shaping dendritic plasticity and neuronal output. Frontiers in Cellular Neuroscience, 18, 1513602. 10.3389/fncel.2024.1513602

Czurkó, A., Huxter, J., Li, Y., Hangya, B., & Muller, R. U. (2011). Theta Phase Classification of Interneurons in the Hippocampal Formation of Freely Moving Rats. Journal of Neuroscience, 31(8), 2938–2947. 10.1523/JNEUROSCI.5037-10.2011

Dimidschstein, J., Chen, Q., Tremblay, R., Rogers, S. L., Saldi, G.-A., Guo, L., Xu, Q., Liu, R., Lu, C., Chu, J., Grimley, J. S., Krostag, A.-R., Kaykas, A., Avery, M. C., Rashid, M. S., Baek, M., Jacob, A. L., Smith, G. B., Wilson, D. E., … Fishell, G. (2016). A viral strategy for targeting and manipulating interneurons across vertebrate species. Nature Neuroscience, 19(12), 1743–1749. 10.1038/nn.4430

Dunot, J., Ribera, A., Pousinha, P. A., & Marie, H. (2023). Spatiotemporal insights of APP function. Current Opinion in Neurobiology, 82, 102754. 10.1016/j.conb.2023.102754

Dupré la Tour, T., Tallot, L., Grabot, L., Doyère, V., van Wassenhove, V., Grenier, Y., & Gramfort, A. (2017). Non-linear auto-regressive models for cross-frequency coupling in neural time series. PLoS Computational Biology, 13(12), e1005893. 10.1371/journal.pcbi.1005893

Ebrahimi, R., Mohammad Soltani, S., Masouri, M. M., Seifi, M., Ghafourian, K., & Noori, S. (2025). Glutamatergic and GABAergic metabolite levels in Alzheimer’s disease: A systematic review and meta-analysis. BMC Neurology, 25, 344. 10.1186/s12883-025-04375-2

Eichenbaum, H. (2017). Prefrontal–hippocampal interactions in episodic memory. Nature Reviews Neuroscience, 18(9), 547–558. 10.1038/nrn.2017.74

Etter, G., van der Veldt, S., Manseau, F., Zarrinkoub, I., Trillaud-Doppia, E., & Williams, S. (2019). Optogenetic gamma stimulation rescues memory impairments in an Alzheimer’s disease mouse model. Nature Communications, 10(1), 5322. 10.1038/s41467-019-13260-9

Fanutza, T., Del Prete, D., Ford, M. J., Castillo, P. E., & D’Adamio, L. (2015). APP and APLP2 interact with the synaptic release machinery and facilitate transmitter release at hippocampal synapses. eLife, 4, e09743. 10.7554/eLife.09743

Fernández-Ruiz, A., Oliva, A., Nagy, G. A., Maurer, A. P., Berényi, A., & Buzsáki, G. (2017). Entorhinal–CA3 dual-input control of spike timing in the hippocampus by theta-gamma coupling. Neuron, 93(5), 1213–1226.e5. 10.1016/j.neuron.2017.02.017

Frere, S., & Slutsky, I. (2018). Alzheimer’s Disease: From Firing Instability to Homeostasis Network Collapse. Neuron, 97(1), 32–58. 10.1016/j.neuron.2017.11.028

Freund, T. F., & Katona, I. (2007). Perisomatic Inhibition. Neuron, 56(1), 33–42. 10.1016/j.neuron.2007.09.012

Gabriele, R. M. C., Abel, E., Fox, N. C., Wray, S., & Arber, C. (2022). Knockdown of Amyloid Precursor Protein: Biological Consequences and Clinical Opportunities. Frontiers in Neuroscience, 16, 835645. 10.3389/fnins.2022.835645

Galarreta, M., & Hestrin, S. (2001). Spike transmission and synchrony detection in networks of GABAergic interneurons. Science, 292(5525), 2295–2299. 10.1126/science.1061395

Garcia, S., Guarino, D., Jaillet, F., Jennings, T. R., Pröpper, R., Rautenberg, P. L., Rodgers, C., Sobolev, A., Wachtler, T., Yger, P., & Davison, A. P. (2014). Neo: An object model for handling electrophysiology data in multiple formats. Frontiers in Neuroinformatics, 8. 10.3389/fninf.2014.00010

Gloveli, T., Dugladze, T., Saha, S., Monyer, H., Heinemann, U., Traub, R. D., Whittington, M. A., & Buhl, E. H. (2005). Differential involvement of oriens/pyramidale interneurones in hippocampal network oscillations in vitro. The Journal of Physiology, 562(Pt 1), 131–147. 10.1113/jphysiol.2004.073007

Goettemoeller, A. M., Banks, E., Kumar, P., Olah, V. J., McCann, K. E., South, K., Ramelow, C. C., Eaton, A., Duong, D. M., Seyfried, N. T., Weinshenker, D., Rangaraju, S., & Rowan, M. J. M. (2024). Entorhinal cortex vulnerability to human APP expression promotes hyperexcitability and tau pathology. Nature Communications, 15(1), 7918. 10.1038/s41467-024-52297-3

Govindpani, K., Calvo-Flores Guzmán, B., Vinnakota, C., Waldvogel, H. J., Faull, R. L., & Kwakowsky, A. (2017). Towards a Better Understanding of GABAergic Remodeling in Alzheimer’s Disease. International Journal of Molecular Sciences, 18(8), 1813. 10.3390/ijms18081813

Hick, M., Herrmann, U., Weyer, S. W., Mallm, J.-P., Tschäpe, J.-A., Borgers, M., Mercken, M., Roth, F. C., Draguhn, A., Slomianka, L., Wolfer, D. P., Korte, M., & Müller, U. C. (2015). Acute function of secreted amyloid precursor protein fragment APPsα in synaptic plasticity. Acta Neuropathologica, 129(1), 21–37. 10.1007/s00401-014-1368-x

Hijazi, S., Heistek, T. S., Scheltens, P., Neumann, U., Shimshek, D. R., Mansvelder, H. D., Smit, A. B., & van Kesteren, R. E. (2020). Early restoration of parvalbumin interneuron activity prevents memory loss and network hyperexcitability in a mouse model of Alzheimer’s disease. Molecular Psychiatry, 25(12), 3380–3398. 10.1038/s41380-019-0483-4

Hijazi, S., Smit, A. B., & van Kesteren, R. E. (2023). Fast-spiking parvalbumin-positive interneurons in brain physiology and Alzheimer’s disease. Molecular Psychiatry, 28(12), 4954–4967. 10.1038/s41380-023-02168-y

Hu, H., Martina, M., & Jonas, P. (2010). Dendritic Mechanisms Underlying Rapid Synaptic Activation of Fast-Spiking Hippocampal Interneurons. Science, 327(5961), 52–58. 10.1126/science.1177876

Jiménez-Balado, J., & Eich, T. S. (2021). GABAergic dysfunction, neural network hyperactivity and memory impairments in human aging and Alzheimer’s disease. Seminars in Cell & Developmental Biology, Special Issue: Myelin Edited by Gonçalo Castelo-Branco and Roman Chrast / Special Issue: Aging in the Nervous System Edited by Mara Mather, 116, 146–159. 10.1016/j.semcdb.2021.01.005

Klausberger, T., & Somogyi, P. (2008). Neuronal Diversity and Temporal Dynamics: The Unity of Hippocampal Circuit Operations. Science, 321(5885), 53–57. 10.1126/science.1149381

Kohus, Z., Káli, S., Rovira-Esteban, L., Schlingloff, D., Papp, O., Freund, T. F., Hájos, N., & Gulyás, A. I. (2016). Properties and dynamics of inhibitory synaptic communication within the CA3 microcircuits of pyramidal cells and interneurons expressing parvalbumin or cholecystokinin. The Journal of Physiology, 594(13), 3745–3774. 10.1113/JP272231

Korte, M., Herrmann, U., Zhang, X., & Draguhn, A. (2012). The role of APP and APLP for synaptic transmission, plasticity, and network function: Lessons from genetic mouse models. Experimental Brain Research, 217(3), 435– 440. 10.1007/s00221-011-2894-6

Kubota, Y., Karube, F., Nomura, M., & Kawaguchi, Y. (2016). The Diversity of Cortical Inhibitory Synapses. Frontiers in Neural Circuits, 10. 10.3389/fncir.2016.00027

Kubota, Y., Kondo, S., Nomura, M., Hatada, S., Yamaguchi, N., Mohamed, A. A., Karube, F., Lübke, J., & Kawaguchi, Y. (2015). Functional effects of distinct innervation styles of pyramidal cells by fast spiking cortical interneurons. eLife, 4, e07919. 10.7554/eLife.07919

Kurucu, H., Colom-Cadena, M., Davies, C., Wilkins, L., King, D., Rose, J., Tzioras, M., Tulloch, J. H., Smith, C., & Spires-Jones, T. L. (2022). Inhibitory synapse loss and accumulation of amyloid beta in inhibitory presynaptic terminals in Alzheimer’s disease. European Journal of Neurology, 29(5), 1311–1323. 10.1111/ene.15043

Lanctot, K. L., Herrmann, N., Rothenburg, L., & Eryavec, G. (2007). Behavioral correlates of GABAergic disruption in Alzheimer’s disease. International Psychogeriatrics, 19(1), 151–158. 10.1017/S1041610206003899

Lasztóczi, B., Tukker, J. J., Somogyi, P., & Klausberger, T. (2011). Terminal field and firing selectivity of cholecystokinin-expressing interneurons in the hippocampal CA3 area. The Journal of Neuroscience: The Official Journal of the Society for Neuroscience, 31(49), 18073–18093. 10.1523/JNEUROSCI.3573-11.2011

Lawrence, J. J., Grinspan, Z. M., & McBain, C. J. (2004). Quantal transmission at mossy fibre targets in the CA3 region of the rat hippocampus. The Journal of Physiology, 554(Pt 1), 175–193. 10.1113/jphysiol.2003.049551

Lawrence, J. J., & McBain, C. J. (2003). Interneuron diversity series: Containing the detonation--feedforward inhibition in the CA3 hippocampus. Trends in Neurosciences, 26(11), 631–640. 10.1016/j.tins.2003.09.007

Lee, S. H., Kang, J., Zhang, C., Bolshakov, V. Y., & Shen, J. (2026). APP family in inhibitory neurons controls inhibitory recruitment and short-term plasticity in the hippocampus. Molecular Brain, 19(1), 35. 10.1186/s13041-026-01296-0

Lopes-dos-Santos, V., van de Ven, G. M., Morley, A., Trouche, S., Campo-Urriza, N., & Dupret, D. (2018). Parsing Hippocampal Theta Oscillations by Nested Spectral Components during Spatial Exploration and Memory-Guided Behavior. Neuron, 100(4), 940–952.e7. 10.1016/j.neuron.2018.09.031

Mallm, J., Tschäpe, J., Hick, M., Filippov, M. A., & Müller, U. C. (2010). Generation of conditional null alleles for *APP* and *APLP2*. Genesis, 48(3), 200–206. 10.1002/dvg.20601

Marchal, C., & Mulle, C. (2004). Postnatal maturation of mossy fibre excitatory transmission in mouse CA3 pyramidal cells: A potential role for kainate receptors. The Journal of Physiology, 561(1), 27–37. 10.1113/jphysiol.2004.069922

Marneffe, C., Moreira-de-Sá, A., Lecomte, S., Erhardt, A., & Mulle, C. (2025). Short term plasticity at hippocampal mossy fiber synapses. Neuroscience, 578, 105–117. 10.1016/j.neuroscience.2024.09.044

Mehr, A., Hick, M., Ludewig, S., Müller, M., Herrmann, U., von Engelhardt, J., Wolfer, D. P., Korte, M., & Müller, U. C. (2020). Lack of APP and APLP2 in GABAergic Forebrain Neurons Impairs Synaptic Plasticity and Cognition. Cerebral Cortex, 30(7), 4044–4063. 10.1093/cercor/bhaa025

Miles, R., Tóth, K., Gulyás, A. I., Hájos, N., & Freund, T. F. (1996). Differences between Somatic and Dendritic Inhibition in the Hippocampus. Neuron, 16(4), 815–823. 10.1016/S0896-6273(00)80101-4

Müller, U. C., Deller, T., & Korte, M. (2017). Not just amyloid: Physiological functions of the amyloid precursor protein family. Nature Reviews Neuroscience, 18(5), 281–298. 10.1038/nrn.2017.29

Olah, V. J., Goettemoeller, A. M., Rayaprolu, S., Dammer, E. B., Seyfried, N. T., Rangaraju, S., Dimidschstein, J., & Rowan, M. J. (2022). Biophysical Kv3 channel alterations dampen excitability of cortical PV interneurons and contribute to network hyperexcitability in early Alzheimer’s. eLife, 11, e75316. 10.7554/eLife.75316

Pachitariu, M., Sridhar, S., Pennington, J., & Stringer, C. (2024). Spike sorting with Kilosort4. Nature Methods, 21(5), 914–921. 10.1038/s41592-024-02232-7

Palop, J. J., Chin, J., Roberson, E. D., Wang, J., Thwin, M. T., Bien-Ly, N., Yoo, J., Ho, K. O., Yu, G.-Q., Kreitzer, A., Finkbeiner, S., Noebels, J. L., & Mucke, L. (2007). Aberrant Excitatory Neuronal Activity and Compensatory Remodeling of Inhibitory Hippocampal Circuits in Mouse Models of Alzheimer’s Disease. Neuron, 55(5), 697–711. 10.1016/j.neuron.2007.07.025

Palop, J. J., & Mucke, L. (2010). Amyloid-β–induced neuronal dysfunction in Alzheimer’s disease: From synapses toward neural networks. Nature Neuroscience, 13(7), 812–818. 10.1038/nn.2583

Palop, J. J., & Mucke, L. (2016). Network abnormalities and interneuron dysfunction in Alzheimer disease. Nature Reviews Neuroscience, 17(12), 777–792. 10.1038/nrn.2016.141

Park, K., Lee, J., Jang, H. J., Richards, B. A., Kohl, M. M., & Kwag, J. (2020). Optogenetic activation of parvalbumin and somatostatin interneurons selectively restores theta-nested gamma oscillations and oscillation-induced spike timing-dependent long-term potentiation impaired by amyloid β oligomers. BMC Biology, 18(1), 7. 10.1186/s12915-019-0732-7

Pelkey, K. A., Chittajallu, R., Craig, M. T., Tricoire, L., Wester, J. C., & McBain, C. J. (2017). Hippocampal GABAergic Inhibitory Interneurons. Physiological Reviews, 97(4), 1619–1747. 10.1152/physrev.00007.2017

Petrache, A. L., Rajulawalla, A., Shi, A., Wetzel, A., Saito, T., Saido, T. C., Harvey, K., & Ali, A. B. (2019). Aberrant Excitatory-Inhibitory Synaptic Mechanisms in Entorhinal Cortex Microcircuits During the Pathogenesis of Alzheimer’s Disease. Cerebral Cortex, 29(4), 1834–1850. 10.1093/cercor/bhz016

Rebola, N., Carta, M., & Mulle, C. (2017). Operation and plasticity of hippocampal CA3 circuits: Implications for memory encoding. Nature Reviews Neuroscience, 18(4), 208–220. 10.1038/nrn.2017.10

Rothman, J. S., & Silver, R. A. (2018). NeuroMatic: An Integrated Open-Source Software Toolkit for Acquisition, Analysis and Simulation of Electrophysiological Data. Frontiers in Neuroinformatics, 12, 14. 10.3389/fninf.2018.00014

Royer, S., Zemelman, B. V., Losonczy, A., Kim, J., Chance, F., Magee, J. C., & Buzsáki, G. (2012). Control of timing, rate and bursts of hippocampal place cells by dendritic and somatic inhibition. Nature Neuroscience, 15(5), 769– 775. 10.1038/nn.3077

Rozov, A., Jappy, D. J., Maltseva, K., Vazetdinova, A., & Valiullina-Rakhmatullina, F. (2024). Dialectics of perisomatic inhibition—The unity and conflict of opposites. Frontiers in Neural Circuits, 18, 1494300. 10.3389/fncir.2024.1494300

Salin, P. A., Scanziani, M., Malenka, R. C., & Nicoll, R. A. (1996). Distinct short-term plasticity at two excitatory synapses in the hippocampus. Proceedings of the National Academy of Sciences of the United States of America, 93(23), 13304–13309. 10.1073/pnas.93.23.13304

Schmid, L. C., Mittag, M., Poll, S., Steffen, J., Wagner, J., Geis, H.-R., Schwarz, I., Schmidt, B., Schwarz, M. K., Remy, S., & Fuhrmann, M. (2016). Dysfunction of Somatostatin-Positive Interneurons Associated with Memory Deficits in an Alzheimer’s Disease Model. Neuron, 92(1), 114–125. 10.1016/j.neuron.2016.08.034

Seabrook, G. R., Smith, D. W., Bowery, B. J., Easter, A., Reynolds, T., Fitzjohn, S. M., Morton, R. A., Zheng, H., Dawson, G. R., Sirinathsinghji, D. J. S., Davies, C. H., Collingridge, G. L., & Hill, R. G. (1999). Mechanisms contributing to the deficits in hippocampal synaptic plasticity in mice lacking amyloid precursor protein. Neuropharmacology, 38(3), 349–359. 10.1016/S0028-3908(98)00204-4

Selkoe, D. J. (2002). Alzheimer’s Disease Is a Synaptic Failure. *Science*, New Series, 298(5594), 789–791. http://www.jstor.org/stable/3832648

Senzai, Y., & Buzsáki, G. (2017). Physiological Properties and Behavioral Correlates of Hippocampal Granule Cells and Mossy Cells. Neuron, 93(3), 691–704.e5. 10.1016/j.neuron.2016.12.011

Sukenik, N., Vinogradov, O., Weinreb, E., Segal, M., Levina, A., & Moses, E. (2021). Neuronal circuits overcome imbalance in excitation and inhibition by adjusting connection numbers. Proceedings of the National Academy of Sciences, 118(12), e2018459118. 10.1073/pnas.2018459118

Szabadics, J., & Soltesz, I. (2009). Functional Specificity of Mossy Fiber Innervation of GABAergic Cells in the Hippocampus. Journal of Neuroscience, 29(13), 4239–4251. 10.1523/JNEUROSCI.5390-08.2009

Topolnik, L., & Tamboli, S. (2022). The role of inhibitory circuits in hippocampal memory processing. Nature Reviews. Neuroscience, 23(8), 476–492. 10.1038/s41583-022-00599-0

Tort, A. B. L., Komorowski, R., Eichenbaum, H., & Kopell, N. (2010). Measuring phase-amplitude coupling between neuronal oscillations of different frequencies. Journal of Neurophysiology, 104(2), 1195–1210. 10.1152/jn.00106.2010

Tukker, J. J., Lasztóczi, B., Katona, L., Roberts, J. D. B., Pissadaki, E. K., Dalezios, Y., Márton, L., Zhang, L., Klausberger, T., & Somogyi, P. (2013). Distinct Dendritic Arborization and In Vivo Firing Patterns of Parvalbumin-Expressing Basket Cells in the Hippocampal Area CA3. Journal of Neuroscience, 33(16), 6809–6825. 10.1523/JNEUROSCI.5052-12.2013

Tzilivaki, A., Tukker, J. J., Maier, N., Poirazi, P., Sammons, R. P., & Schmitz, D. (2023). Hippocampal GABAergic interneurons and memory. Neuron, 111(20), 3154–3175. 10.1016/j.neuron.2023.06.016

Vancura, B., Geiller, T., Grosmark, A., Zhao, V., & Losonczy, A. (2023). Inhibitory control of sharp-wave ripple duration during learning in hippocampal recurrent networks. Nature Neuroscience, 26(5), 788–797. 10.1038/s41593-023-01306-7

Varga, C., Golshani, P., & Soltesz, I. (2012). Frequency-invariant temporal ordering of interneuronal discharges during hippocampal oscillations in awake mice. Proceedings of the National Academy of Sciences, 109(40), E2726–E2734. 10.1073/pnas.1210929109

Verret, L., Mann, E. O., Hang, G. B., Barth, A. M. I., Cobos, I., Ho, K., Devidze, N., Masliah, E., Kreitzer, A. C., Mody, I., Mucke, L., & Palop, J. J. (2012). Inhibitory interneuron deficit links altered network activity and cognitive dysfunction in Alzheimer model. Cell, 149(3), 708–721. 10.1016/j.cell.2012.02.046

Viana da Silva, S., Haberl, M. G., Gaur, K., Patel, R., Narayan, G., Ledakis, M., Fu, M. L., de Castro Vieira, M., Koo, E. H., Leutgeb, J. K., & Leutgeb, S. (2023). Localized APP expression results in progressive network dysfunction by disorganizing spike timing. *Neuron*, S0896627323007572. 10.1016/j.neuron.2023.10.001

Vida, I., Bartos, M., & Jonas, P. (2006). Shunting inhibition improves robustness of gamma oscillations in hippocampal interneuron networks by homogenizing firing rates. Neuron, 49(1), 107–117. 10.1016/j.neuron.2005.11.036

Viney, T. J., Lasztoczi, B., Katona, L., Crump, M. G., Tukker, J. J., Klausberger, T., & Somogyi, P. (2013). Network state-dependent inhibition of identified hippocampal CA3 axo-axonic cells in vivo. Nature Neuroscience, 16(12), 1802–1811. 10.1038/nn.3550

Virtanen, P., Gommers, R., Oliphant, T. E., Haberland, M., Reddy, T., Cournapeau, D., Burovski, E., Peterson, P., Weckesser, W., Bright, J., van der Walt, S. J., Brett, M., Wilson, J., Millman, K. J., Mayorov, N., Nelson, A. R. J., Jones, E., Kern, R., Larson, E., … van Mulbregt, P. (2020). SciPy 1.0: Fundamental algorithms for scientific computing in Python. Nature Methods, 17(3), 261–272. 10.1038/s41592-019-0686-2

Wang, B., Wang, Z., Sun, L., Yang, L., Li, H., Cole, A. L., Rodriguez-Rivera, J., Lu, H.-C., & Zheng, H. (2014). The Amyloid Precursor Protein Controls Adult Hippocampal Neurogenesis through GABAergic Interneurons. Journal of Neuroscience, 34(40), 13314–13325. 10.1523/JNEUROSCI.2848-14.2014

Wang, P., Yang, G., Mosier, D. R., Chang, P., Zaidi, T., Gong, Y.-D., Zhao, N.-M., Dominguez, B., Lee, K.-F., Gan, W.-B., & Zheng, H. (2005). Defective neuromuscular synapses in mice lacking amyloid precursor protein (APP) and APP-Like protein 2. The Journal of Neuroscience: The Official Journal of the Society for Neuroscience, 25(5), 1219–1225. 10.1523/JNEUROSCI.4660-04.2005

Wang, X. J., & Buzsáki, G. (1996). Gamma oscillation by synaptic inhibition in a hippocampal interneuronal network model. The Journal of Neuroscience: The Official Journal of the Society for Neuroscience, 16(20), 6402– 6413. 10.1523/JNEUROSCI.16-20-06402.1996

Wester, J. C., & McBain, C. J. (2014). Behavioral state-dependent modulation of distinct interneuron subtypes and consequences for circuit function. *Current Opinion in Neurobiology*, SI: Neuromodulation, 29, 118–125. 10.1016/j.conb.2014.07.007

Wu, Z., Guo, Z., Gearing, M., & Chen, G. (2014). Tonic inhibition in dentate gyrus impairs long-term potentiation and memory in an Alzheimer’s [corrected] disease model. Nature Communications, 5, 4159. 10.1038/ncomms5159

Xu, Y., Zhao, M., Han, Y., & Zhang, H. (2020). GABAergic Inhibitory Interneuron Deficits in Alzheimer’s Disease: Implications for Treatment. Frontiers in Neuroscience, 14, 660. 10.3389/fnins.2020.00660

Zott, B., Busche, M. A., Sperling, R. A., & Konnerth, A. (2018). What Happens with the Circuit in Alzheimer’s Disease in Mice and Humans? Annual Review of Neuroscience, 41(1), 277–297. 10.1146/annurev-neuro-080317-061725

Zott, B., Simon, M. M., Hong, W., Unger, F., Chen-Engerer, H.-J., Frosch, M. P., Sakmann, B., Walsh, D. M., & Konnerth, A. (2019). A vicious cycle of β amyloid–dependent neuronal hyperactivation. Science, 365(6453), 559– 565. 10.1126/science.aay0198

