## supplementary figures for "Implication of the amyloid precursor protein expressed in GABAergic interneurons for CA3 neuronal dynamics"

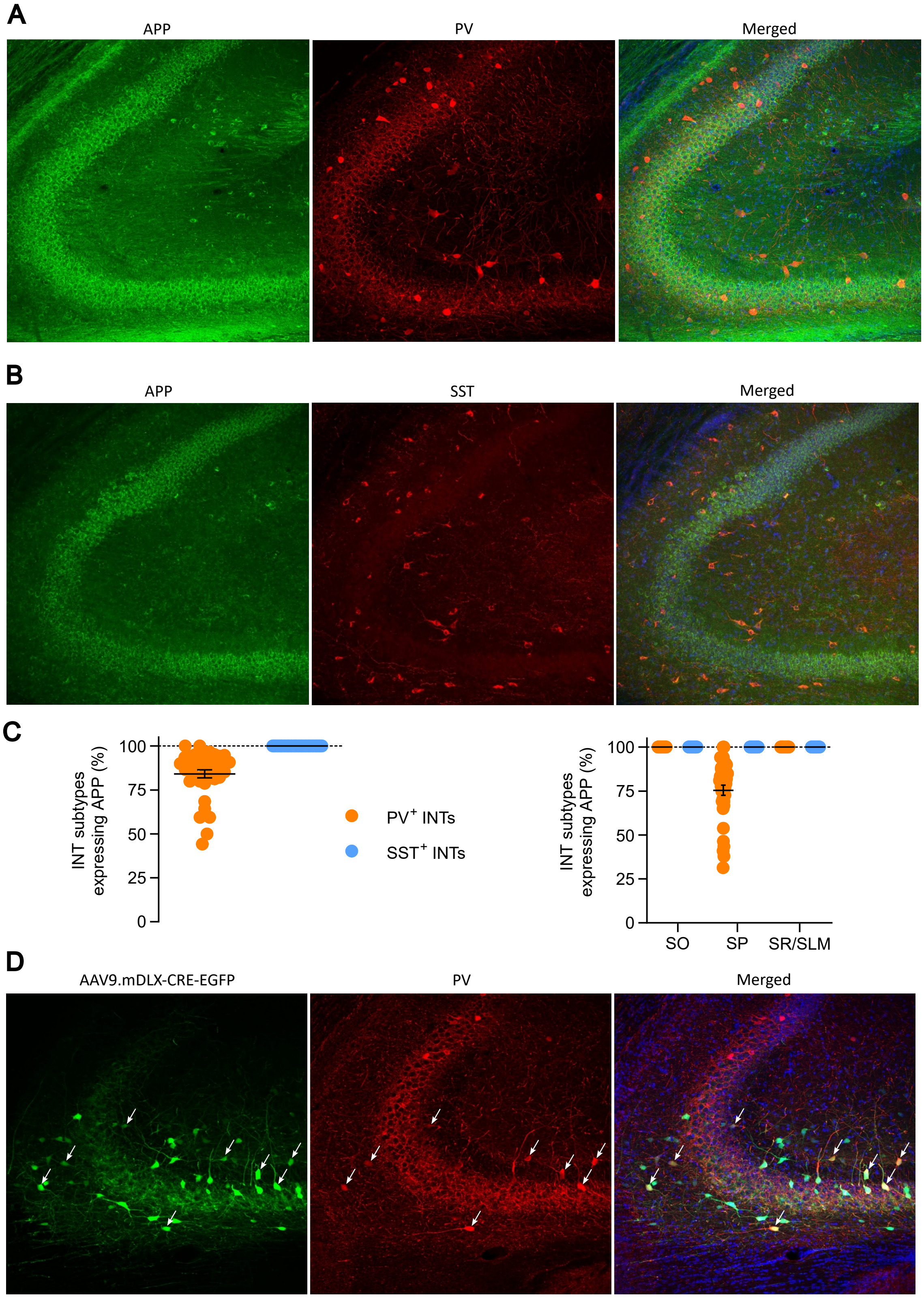


**Supplementary figure 1**

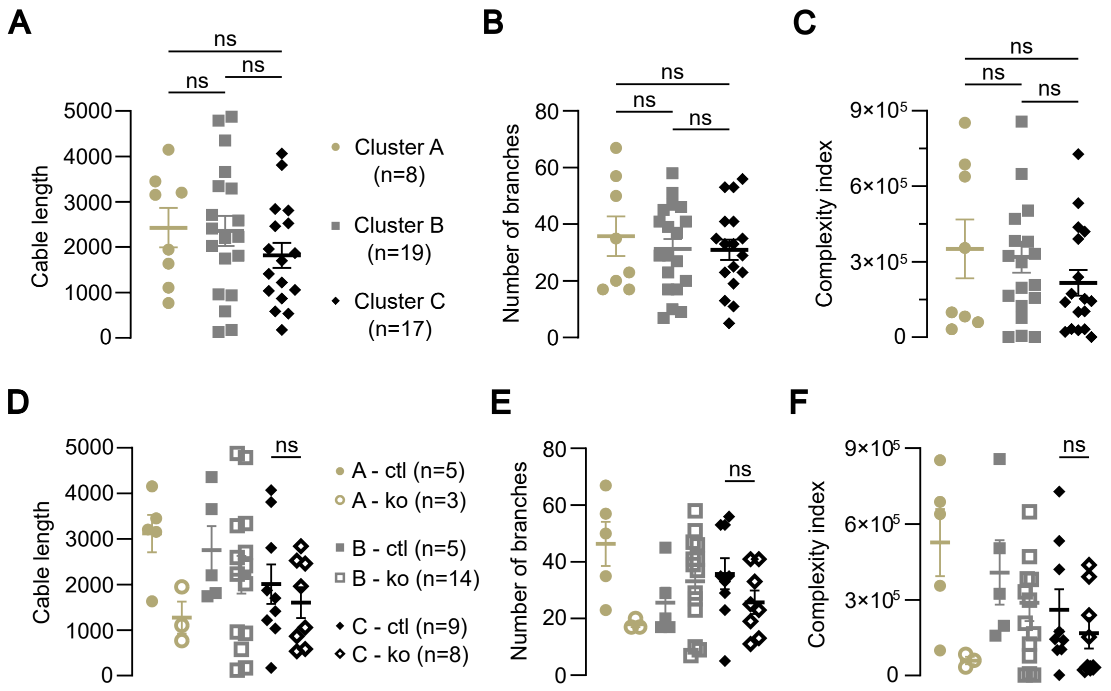


**Supplementary figure 2**

**A**

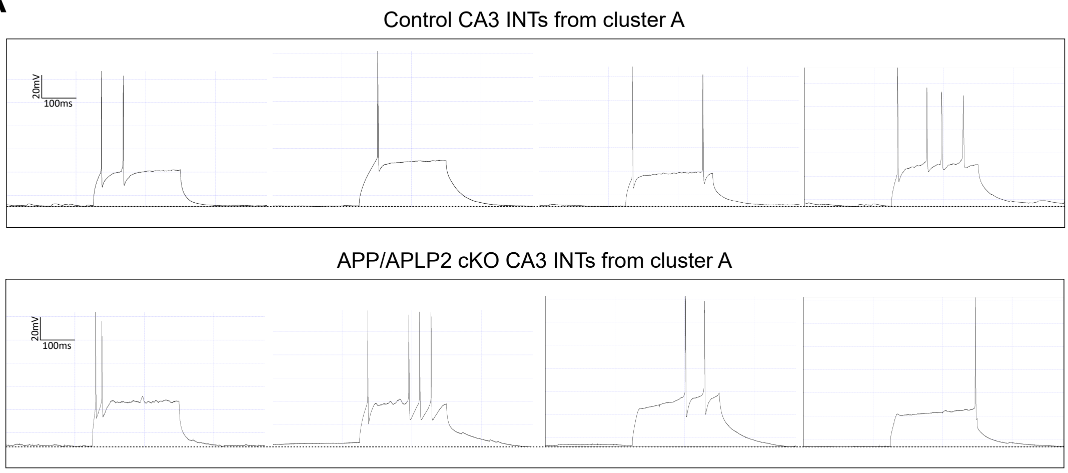


**B**

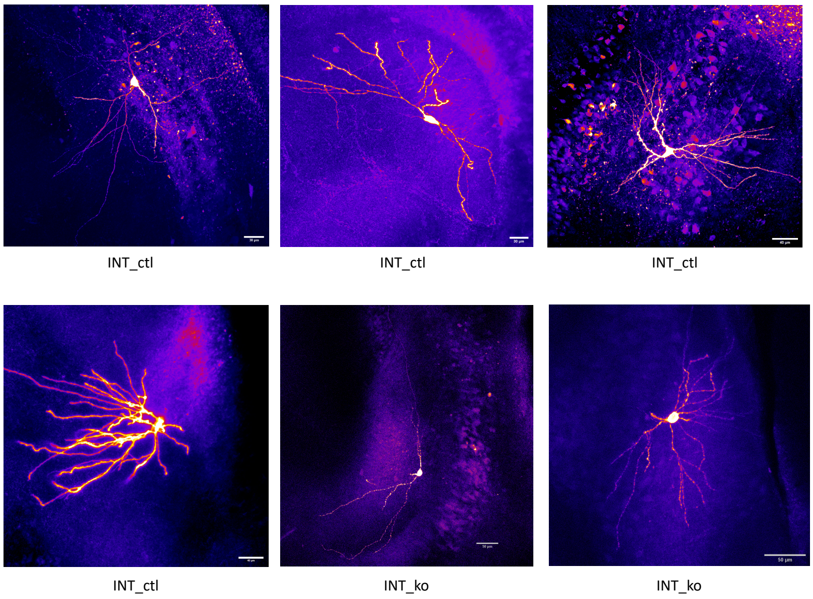


**Supplementary figure 3**

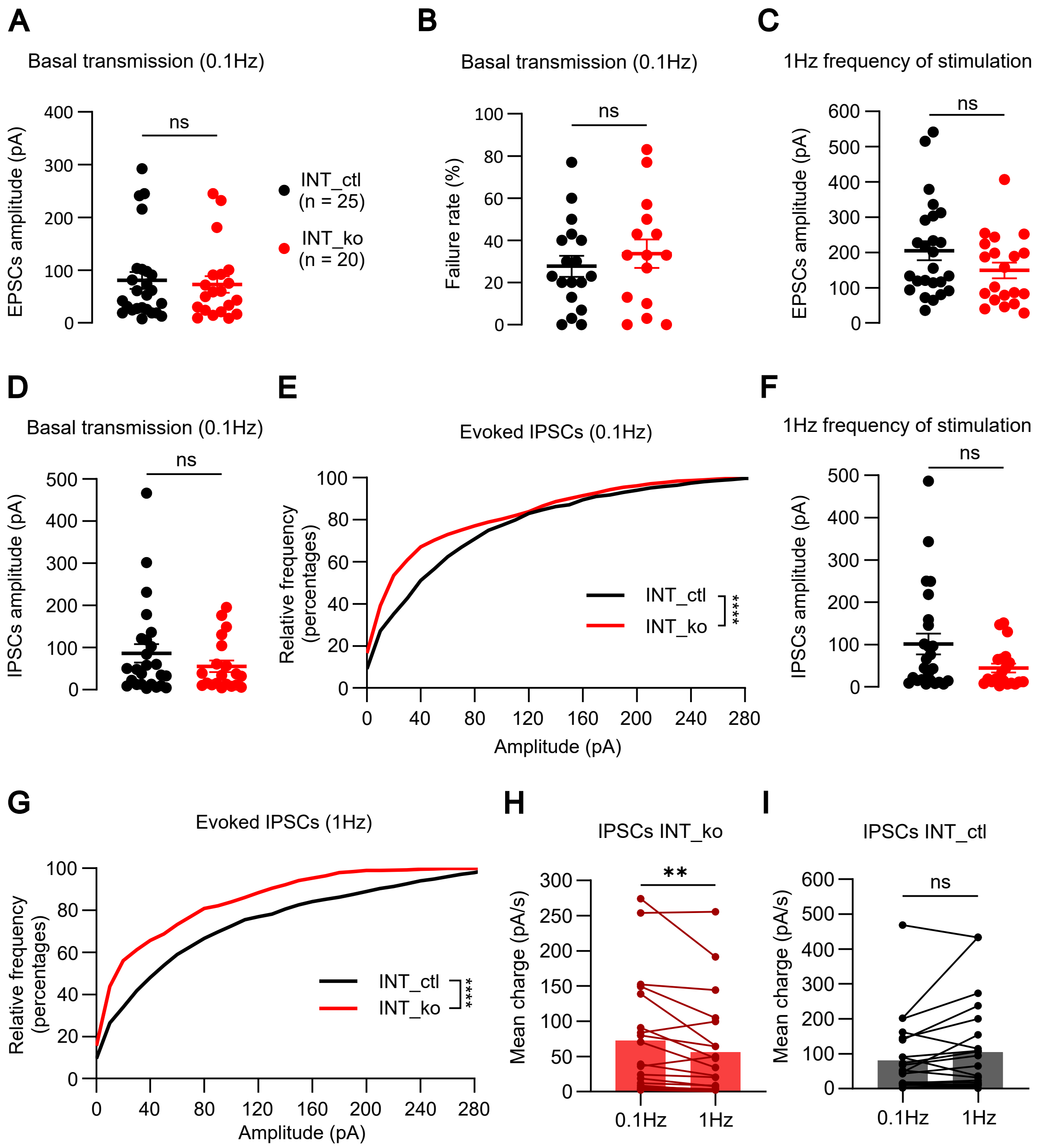


**Supplementary figure 4**

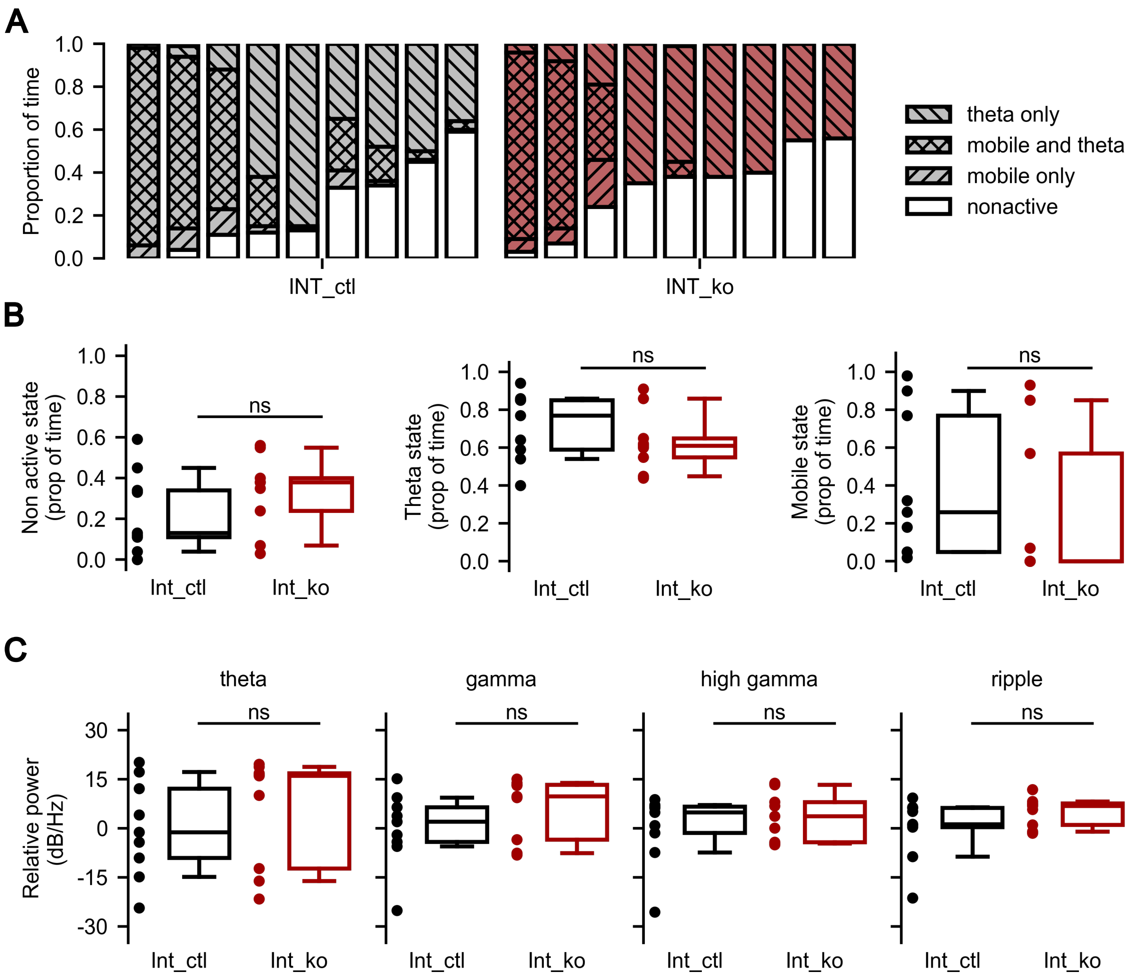


**Supplementary figure 5**

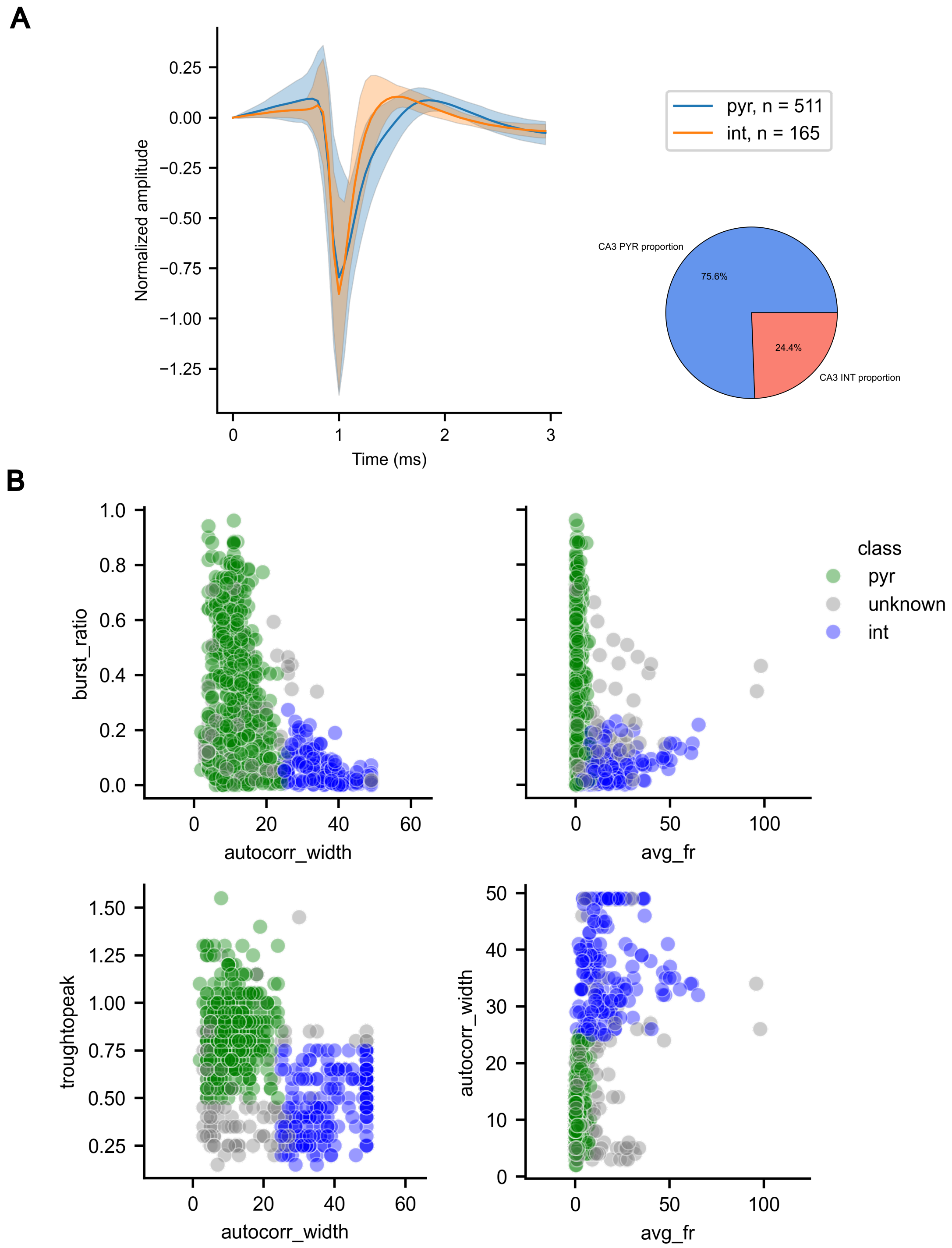


**Supplementary figure 6**

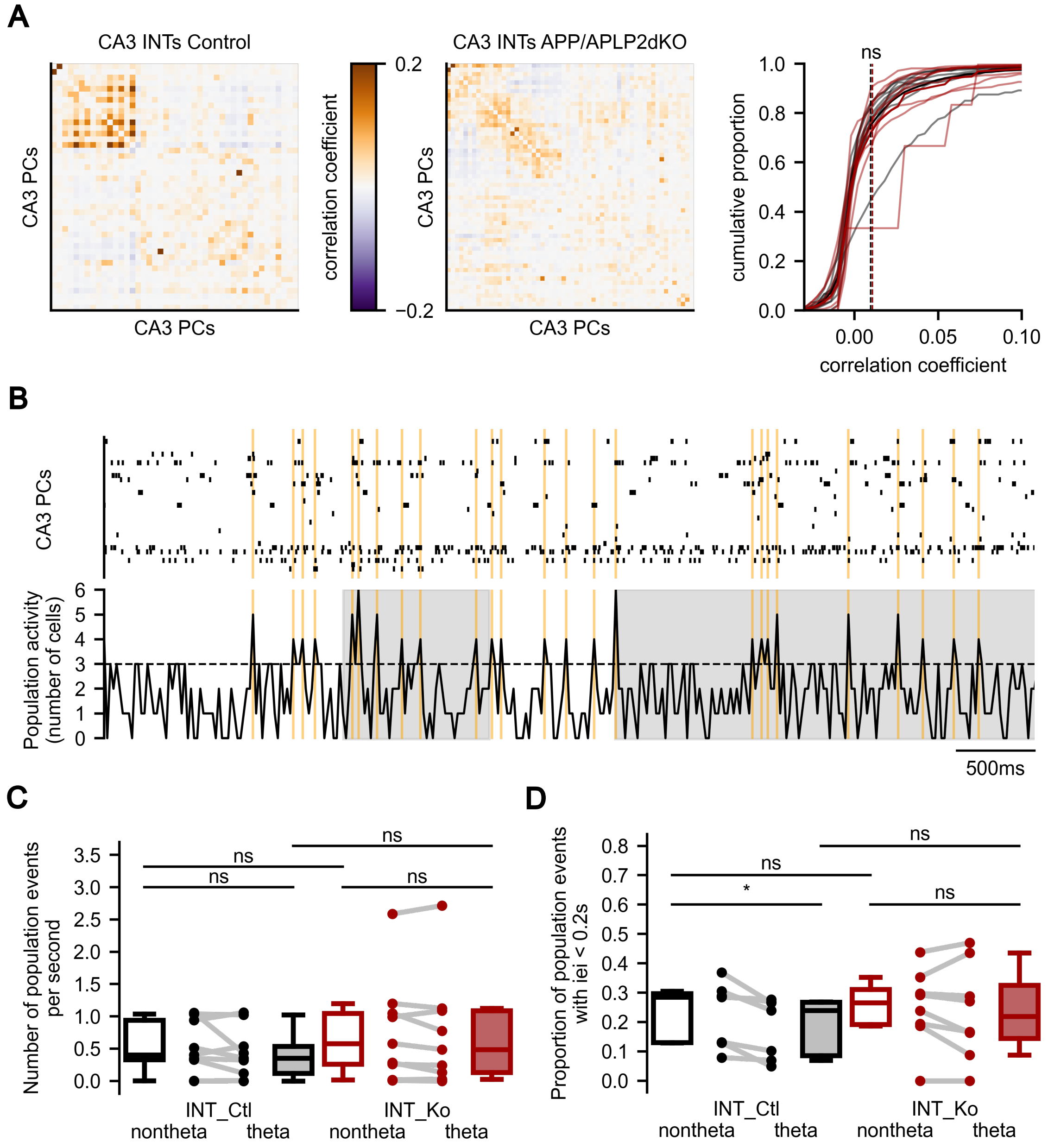


**Supplementary figure 7**
